# Black-box testing in motor sequence learning

**DOI:** 10.1101/2021.12.01.470563

**Authors:** Pablo Maceira-Elvira, Jan E. Timmermann, Traian Popa, Anne-Christine Schmid, John W. Krakauer, Takuya Morishita, Maximilian J. Wessel, Friedhelm C. Hummel

## Abstract

During learning of novel motor sequences, practice leads to the consolidation of hierarchical structures, namely motor chunks, facilitating the accurate execution of sequences at increasing speeds. Recent studies show that such hierarchical structures are largely represented upstream of the primary motor cortex in the motor network, suggesting their function to be more related to the encoding, storage, and retrieval of sequences rather than their sole execution. We isolated different components of motor skill acquisition related to the consolidation of spatiotemporal features and followed their evolution over training. We found that optimal motor skill acquisition relies on the storage of the spatial features of the sequence in memory, followed by the optimization of its execution and increased execution speeds (i.e., a shift in the speed-accuracy trade-off) early in training, supporting the model proposed by Hikosaka in 1999. Contrasting the dynamics of these components during ageing, we identified less-than-optimal mechanisms in older adults explaining the observed differences in performance. We applied noninvasive brain stimulation in an attempt to support the aging brain to compensate for these deficits. The present study found that anodal direct current stimulation applied over the motor cortex restored the mechanisms involved in the consolidation of spatial features, without directly affecting the speed of execution of the sequence. This led older adults to sharply improve their accuracy, resulting in an earlier yet gradual emergence of motor chunks. The results suggest the early storage of the sequence in memory, largely independent of motor practice, is crucial for an optimal motor acquisition and retrieval of this motor behavior. Nevertheless, the consolidation of optimal temporal patterns, detected as motor chunks at a behavioral level, is not a direct consequence of storing the sequence elements, but rather of motor practice.

## Introduction

Completing daily life activities often requires the sequential execution of actions in a specific order. A large amount of research has focused on how humans acquire sequential motor skills using well established experimental paradigms alongside different imaging techniques to study the processes that lead to skill improvement ^1^. One of these paradigms, known as the sequential finger-tapping task ^2,3^, has been used in past years due to its similarity to certain activities requiring higher dexterous skill, such as piano playing or typing on a computer. Performance improvement of a sequence-tapping task is characterized by a shift in the speed-accuracy tradeoff, in which the speed of execution of the motor sequence increases without sacrificing the accuracy ^4^. The execution of sequential elements at increasing speeds leads to the spontaneous emergence of execution patterns ^5,6^, namely motor chunks ^7^, which reduce mental load ^8^ and facilitate a further increase in speed without sacrificing accuracy ^9^.

Recent discussions about this type of motor task are concerned with its validity for probing changes in motor ability ^10^. Motor chunks seem to be crucial for the optimization of such a task. In spite of the ongoing debate on the role of the primary motor cortex (M1) in motor skill acquisition ^11,12^, recent studies have not found a representation of such structures in the primary motor cortex ^13,14^, so it would appear the task is probing mainly the cognitive aspects of motor learning, specifically the efficient retrieval of the sequence elements from memory (for a detailed discussion, please see ^15^). Nevertheless, most studies looking at the consolidation of motor chunks have been done in healthy young adults, a population in which the involved mechanisms, such as the encoding, storage and the successful retrieval of sequence elements may be acting too quickly to be captured by the applied methods.

Black-box testing (https://en.wikipedia.org/wiki/Black-box_testing), a common software testing technique, examines the functionality of an application by comparing the expected functionality of the system (i.e., requirement) and its actual performance. This approach can be applied to biological systems as well. For example, Shadmehr and Krakauer ^16^ compared computational models describing motor control to specific populations of patients with lesions in the central nervous system, mapping different model parameters to lesioned brain areas and attributing distinct roles to them (e.g., state estimation, optimization, etc.). Similarly, understanding the mechanisms involved in motor sequence learning may be better achieved through the juxtaposition of individuals constituting the requirement (e.g., young adults, depicting optimal performance) and individuals in which the involved mechanisms may no longer function optimally (e.g., older adults).

Previous research shows neurophysiological, structural and functional changes occurring in the aging brain that lead to a decline in cognitive ^17^ and motor functions ^18–22^; for review, please refer to ^23,24^. As such, motor skill acquisition is typically diminished in older adults ^3,25,26^. However, the application of anodal transcranial direct current stimulation (atDCS) to the motor cortex seems to enhance the motor skill acquisition ^3,27^. Even though the mechanisms of action of atDCS in individuals are complex and not yet entirely understood, its application can be used as an additional probe in the “black-box testing” of motor skill acquisition.

We designed a study intended to identify (a) the main factors leading to differences in motor skill acquisition with aging, and (b) the effect of applying noninvasive brain stimulation during motor training. Comparing different components of motor skill acquisition in young and older adults, constituting the extremes of performance in this study, we found that the improvement of the sequence-tapping task is maximized by the early consolidation of the spatial properties of the sequence in memory (i.e., sequence order), leading to a reduced error of execution, and by the optimization of its temporal features (i.e., chunking). We found the consolidation of spatiotemporal features to occur early in training in young adults, suggesting the emergence of motor chunks to be a direct consequence of committing the sequence elements to memory. This process, seemingly less efficient in older adults, could be partially restored using atDCS by enabling the early consolidation of spatial features, allowing them to prioritize the increase of their speed of execution, ultimately leading to an earlier consolidation of motor chunks. This separate consolidation of spatial and temporal features seen in older adults suggests that the emergence of temporal patterns, commonly identified as motor chunks at a behavioral level, stem from the optimization of the execution of the motor sequence resulting from practice, which can occur only after the sequence order has been stored in memory.

## Results

### Age-related behavioral differences in the execution and practice of a sequence-tapping task

We studied differences in motor performance related to healthy aging using a well-established sequence-tapping task ^2,3^, and followed their evolution during training. We recruited a cohort of 52 healthy adults belonging to three age groups: young (18-30 y/o; n = 22, 13 female; age*μ* = 24.7 y/o), middle-aged (50-65 y/o; n = 15, 9 female; age*μ* = 57.4 y/o), and older (>65 y/o; n = 15, 8 female; age*μ* = 74.1 y/o). Each participant trained for twenty minutes each day on five consecutive days. The training consisted of six 90-second training blocks interspaced by 90-second blocks of rest. The participants had to replicate a nine-digit sequence displayed on a screen, as quickly and as accurately as possible, using their left (nondominant) hand. We inserted a seventh block with a different sequence (*i.e.,* “catch” block) halfway through training to evaluate the difference between the pure motor execution of a random sequence and that of the trained sequence. The participants returned on day 10 and day 60, from the beginning of training, to evaluate the long-term retention of the learned sequence.

*Figure 1a* shows the main results of this experiment. We found no transfer of learning from the training sequence to the catch blocks, so we removed these blocks for the subsequent analyses (please refer to *Supplementary Materials* to find the scores including the catch blocks). We scored participants by considering the number of correct sequences produced in each block, weighted by the ratio of correct to absolute number of sequences (*i.e.,* percent correct). To capture individual improvement on the training sequence, we corrected individual scores by subtracting the score in the first block from the scores in the following blocks as a normalization procedure (please refer to *Supplementary Materials* for more information on the choice for scoring, as well as to find the uncentered scores of all groups).

**Figure 1.**
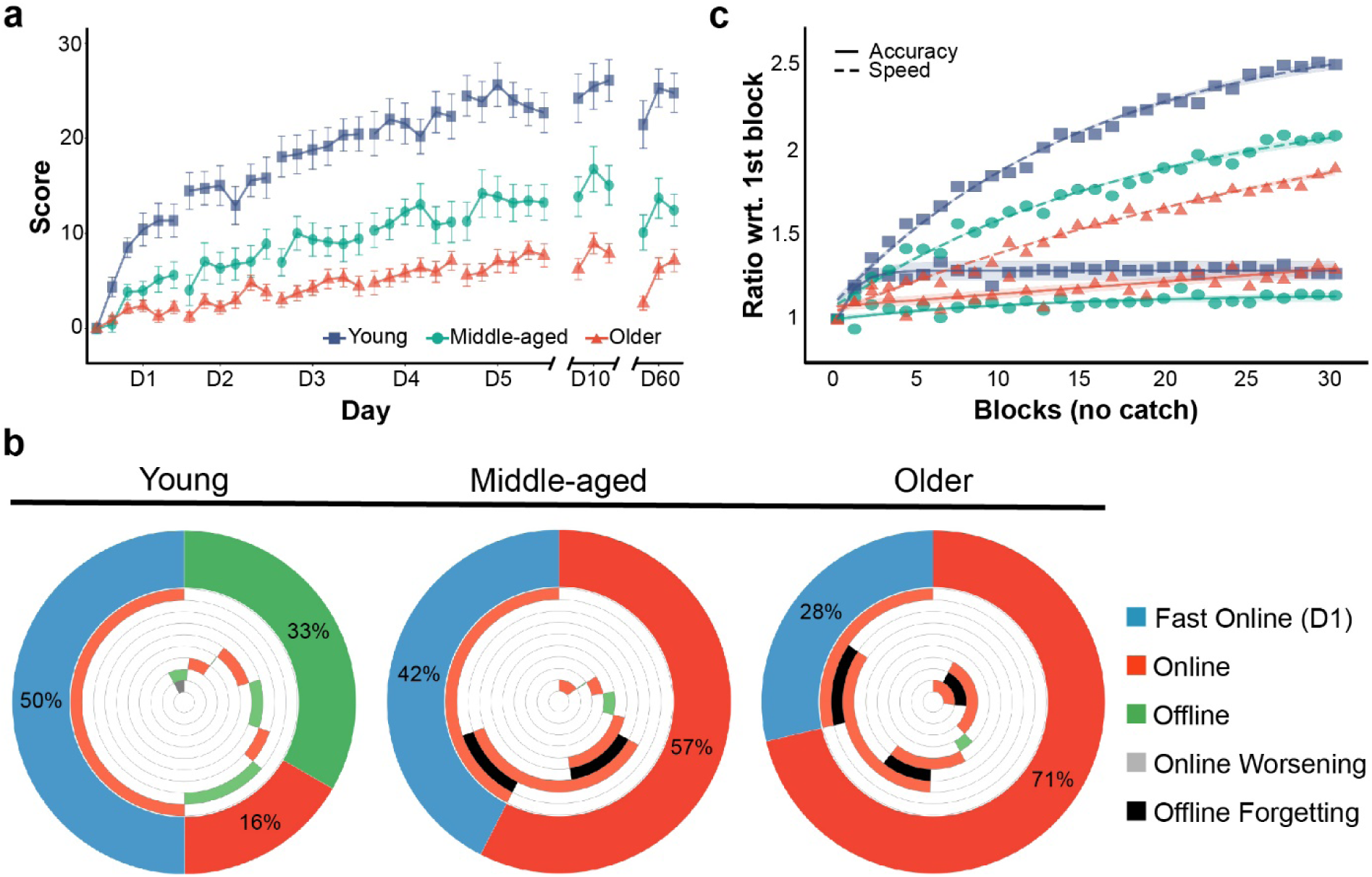
Motor skill acquisition in Experiment 1. **a)** Scores generated during Experiment 1, in which participants trained on the motor sequence with no stimulation. Scores are averaged per age group, and the error bars correspond to the standard error of the mean. The blocks of “catch trials” with a different sequence (one block every training day) are not presented. **b)** Percentage of total learning over the entire training week represented by different aspects of learning (i.e., fast-online learning during D1, online learning during D2-5, and offline learning between training days). The outer ring captures the proportion of total learning by these three aspects, while the inner rings present their time course during the week (anticlockwise): 1^st^ inner circle is the online performance gain during D1, 2^nd^ inner circle is the offline performance gain between D1 and D2, 3^rd^ inner circle is the online gain during D2, etc. Orange and green represent improvements, while black and gray represent worsening of performance. Please note that young adults show offline improvement between days, while middle-aged and older adults not only lack such improvement but also worsen overnight. **c)** Speed and accuracy, normalized to the values in the first block of training, reflect relative changes with respect to initial levels. All groups show consistent increases in speed with similar dynamics; relative differences in magnitude between age groups show young adults being fastest and older adults slowest. Please note the different accuracy dynamics when comparing young adults, who sharply improve accuracy on the first day, to older adults, who gradually improve accuracy during the entire training week. Of particular import is the fact of all age groups displaying consistently increasing speeds, without ever dropping in accuracy, constituting a shift in the speed-accuracy tradeoff. The shading represents the 95% confidence interval for the logarithmic curve fitting (this type of curve is for display purposes only and not included in the LME analysis).

We used a linear mixed-effects (LME, please refer to *Methods* for details) model to quantify differences between groups. Scores on the fifth day (*i.e.,* total learning, relative to the first block of training) were significantly higher in the young adults than in the middle-aged (T_[55]_ = 10.78, d = 2.61, p < 0.0001) and older (T_[55]_ = 17.08, d = 4.14, p < 0.0001) adults, with the middle-aged group scoring significantly higher than the older group (T_[55]_ = 6.3, d = 1.52, p = 0.01). At the follow-up testing days (*i.e.,* day 10 and day 60), the relative differences between the age groups persisted. Performance in all groups continued to increase significantly by the tenth day (T_[414]_ = 1.39, d = 0.31, p = 0.01), but dropped back to the level of day five on day 60.

The performance of individuals executing explicit motor sequence learning tasks has been characterized by nonlinear improvement dynamics, showing sharp improvements occurring during the first training day and modest improvements in subsequent days ^28^. Therefore, we compared the rate of improvement (*i.e.,* slopes) between age groups on each training day. We found a marked difference on the first day, where the slope for the young group was significantly steeper than the slope for the middle-aged (T_[245]_ = 0.88, d = 0.99, p = 0.008) and older (T_[245]_ = 1.59, d = 1.8, p < 0.0001) groups. In young individuals, this slope was significantly steeper than that on the second day (T_[245]_ = 1.72, d = 1.94, p < 0.0001). Differences between slopes in middle-aged and older groups on the first day and differences among all groups from the second day onward were not significant. This suggested that the dynamics of the learning process, especially on the first day, are one of the main factors leading to the differences observed by the end of training.

We also tested overnight consolidation (*i.e.,* offline learning), which is known to be diminished in aging populations ^29,30^ due to different sleep patterns, such as lower quality or fragmented sleep ^31^. We found offline learning to be significantly higher in the young adults than in the middle-aged (T_[196]_ = 2.53, d = 0.55, p = 0.002) and older (T_[196]_ = 2.63, d = 0.57, p = 0.001) adults, with no differences found between the middle-aged and older groups (T_[196]_ = 0.1, d = 0.02, p = 0.99).

*Figure 1b* shows the proportion of total learning represented by fast online learning during the first day, slower online learning during the subsequent days, and offline learning between training days. Of note was the lack of offline learning in the middle-aged and older adults, which was replaced by offline forgetting. Previous research has shown learning consolidation after sleep for finger-tapping tasks ^2,32^, an effect apparent here in young participants. The extent of this consolidation might depend on different sleep-related factors ^33^. In older adults, previous research has shown impaired consolidation of motor learning ^29,30^, potentially related to reduced sleep spindle oscillations and an associated decrease in activity in the corticostriatal network ^34^. Diminished sleep quality in older adults, derived from changes in the circadian rhythm and fragmented sleep ^31^, could also contribute to the lack of offline gains.

### Age-dependent differences in speed and accuracy

As motor skill acquisition refers to the practice-related increase in speed and accuracy in the execution of a motor task ^35^, these parameters could explain differences in the slope on the first day. Speed in the young adults was significantly higher than that in the middle-aged (T_[49]_ = 7.4, d = 2.64, p = 0.0002) and older (T_[49]_ = 12.33, d = 4.40, p < 0.0001) adults, and speed in the middle-aged adults was higher than that in the older adults (T_[49]_ = 4.93, d = 1.76, p = 0.02). Accuracy on the first day was not significantly different between age groups, but the young group was significantly more accurate than the older group on day two (T_[76]_ = 0.07, d = 0.82, p = 0.01) and day three (T_[76]_ = 0.07, d = 0.81, p = 0.01).

We normalized the speed and accuracy in each group to study the dynamics of these two parameters. *Figure 1c* shows the changes in both speed and accuracy relative to the first block of training for all three groups (please refer to *Supplementary Materials* for more details on the calculation of speed and accuracy). Speed consistently increased across training in all age groups, albeit to different extents (*Figure 1c*, dashed lines). Accuracy in the young and older adults followed different dynamics; starting from similar levels of accuracy on the first day, the young participants sharply increased their accuracy in the early stages of training and reached a plateau, whereas the older group gradually reached its maximum accuracy over the course of the training week (*Figure 1c*, solid lines). In other words, young adults improved their execution following a pattern reminiscent of the model presented by Hikosaka and colleagues ^36^, in which the spatial coordinates of the task (i.e., the accurate mapping of numbers to fingers stored in memory) are optimized before the motor coordinates (i.e., rapid execution of motion). In contrast, older adults seem to develop both coordinates in parallel, gradually increasing both speed and accuracy.

### Motor chunks and age-related differences

Motor chunking is a well-established model of how individuals approach sequential tasks ^6^. In the hierarchical model of sequencing, long sequences are segmented into shorter chunks ^37^, which consist of groups of individual movements prepared and buffered for their rapid successive execution, to balance execution efficiency and computational complexity ^38^. We extracted chunking patterns from every participant by applying a cluster-based algorithm (please refer to *Methods* for details) that characterized their strategies for each day with a binary, nine-digit sequence. *Figure 2* depicts a radial visualization of the patterns extracted for each participant on day one.

**Figure 2.**
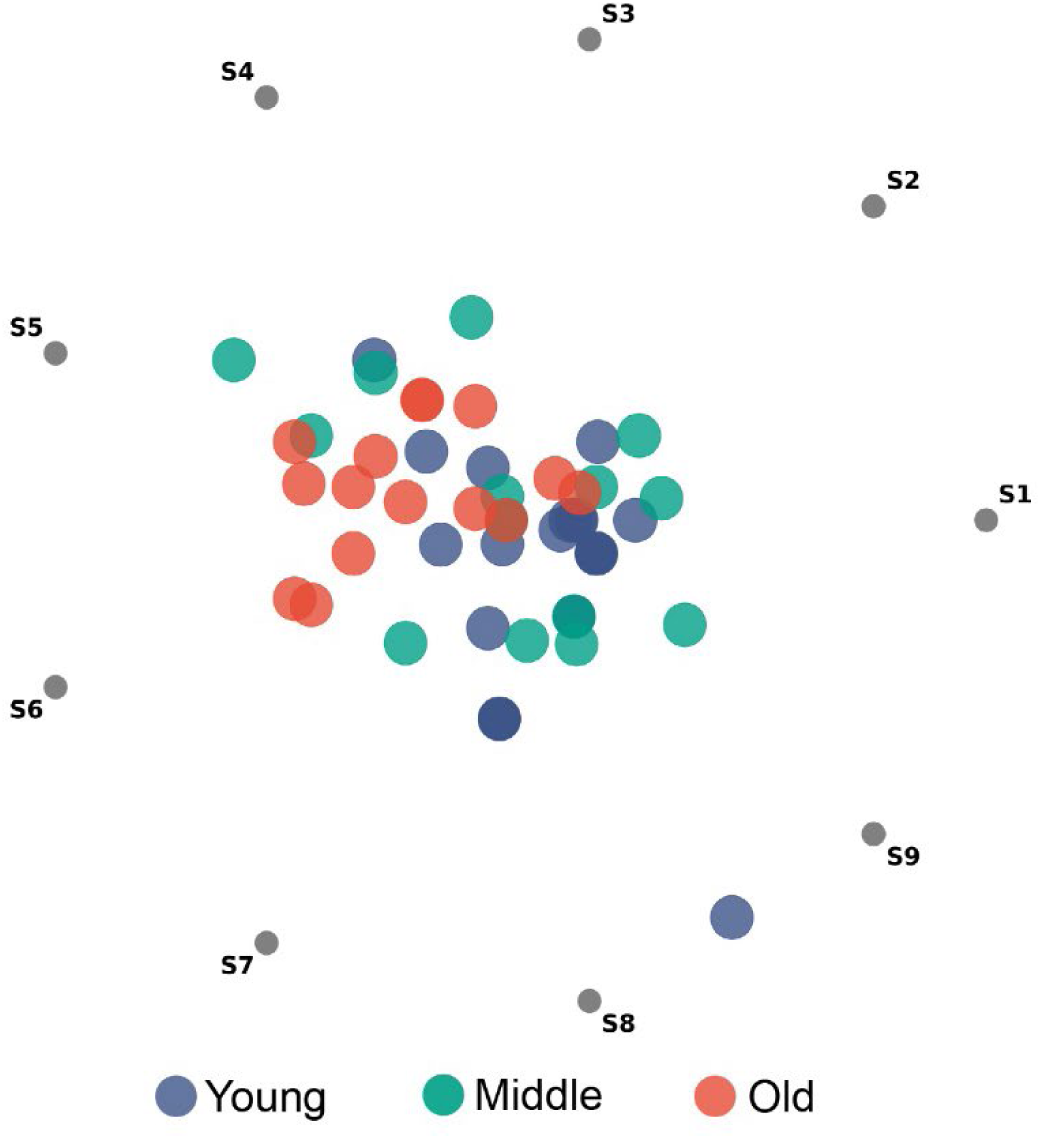
Radial visualization of chunking patterns generated during the first day of training by participants in the first experiment that involved motor training without stimulation. Gray dots on the perimeter of the circle (S1, S2, …, S9) correspond to each bin of the chunking pattern extracted from each participant. Each value of the chunking patterns acts as a “rope” pulling on the data points. For example, a chunking strategy grouping almost exclusively the last three elements of the sequence (labeled [0, 0, 0, 0, 0, 0, 0, 1, 1]) would cause bits S8 and S9 to pull on the data point, resulting in the blue dot situated between S8 and S9 on the plot.

The young and older adults clustered more densely in specific regions, whereas middle-aged adults were distributed between the other two. To quantify the differences in chunking strategies between age groups, we fitted a support vector classifier (SVC) to patterns generated by young and older adults on the first day (please refer to *Methods* for more details), as they represented the extremes in speed and general performance. After fine-tuning the classifier, we extracted the distance of each nine-dimensional data point characterizing the chunking patterns to the decision boundary separating the “young” and “old” classes; we hereafter refer to this parameter as the “chunking distance”. *Figure 3a* shows the extracted chunking distances for the first and last days of training, as well as for day 60 post-training (i.e., the last follow-up session). On the first day, patterns from most young and older participants were correctly classified as such, confirming the presence of the clusters we detected by visual inspection in *Figure 2*. Regarding the middle-aged adults, most seemed to generate patterns that were more similar to those of the young adults on the first day, with some exceptions.

**Figure 3.**
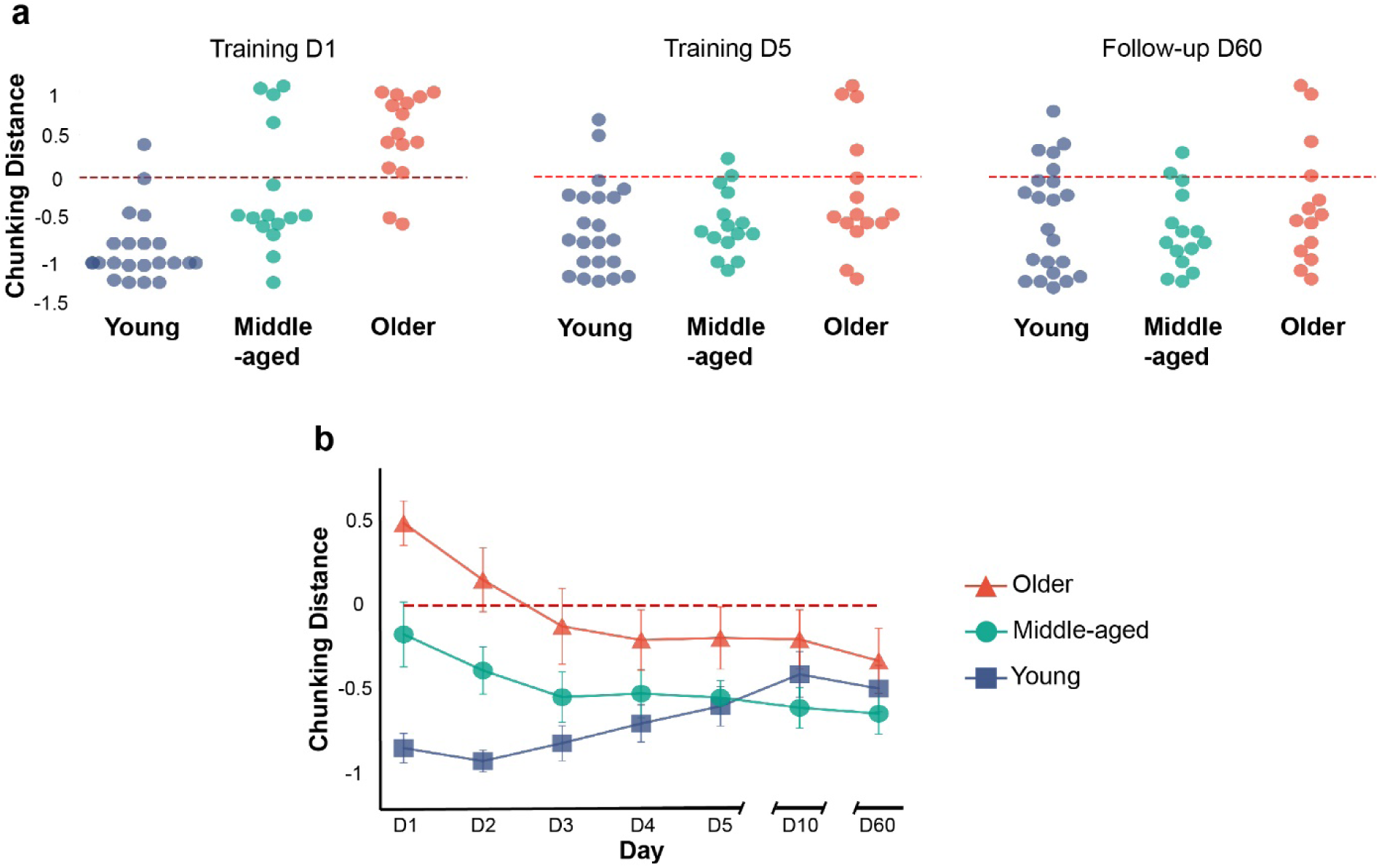
Evolution of chunking patterns during motor skill acquisition in Experiment 1. **a)** Chunking distance for patterns generated on the first and last days of training, as well as on the last follow-up session (i.e., day 60), for each group in Experiment 1. **b)** Average chunking distance for each group on each day, with its corresponding standard error. The red dashed line in both panels indicates the boundary between the two classes that characterizes the chunking strategies of the young and older participants during the first training day, with any distance larger than zero being labeled “old” and any distance smaller than zero labeled “young”.

We used the same model to separate chunking patterns for the remaining days, which consisted of data points previously unseen by the classifier. *Figure 3a* shows that most middle-aged and older adults generated chunking patterns similar to those of the young adults by the end of training, with young adults not significantly changing their strategies. This is consistent with reports from the literature showing “young-like” chunking in older adults after more prolonged training ^39^. *Figure 3b* shows that this process was more gradual in older adults, which was consistent with the gradual increases in accuracy shown in *Figure 1c*.

### Motor training in combination with atDCS in an aging population

We conducted a separate experiment (Experiment 2) following the same design as in Experiment 1, with the addition of atDCS applied over the motor cortex contralateral to the training hand during motor skill acquisition to enhance performance of the task. We recruited a new cohort of 61 healthy adults belonging to the same age groups: young (18-30 y/o; n = 19, 15 female; age*μ* = 24.4), middle-aged (50-65 y/o; n = 19, 11 female; age*μ* = 58), and older (>65 y/o; n = 23, 13 female; age*μ* = 71.2) groups. We randomly assigned participants in each age group to receive either real (*i.e.,* verum; young = 10, middle-aged = 9, older = 14) or placebo stimulation (young = 9, middle-aged = 10, older = 9). The participants trained for the same amount of time as the unstimulated cohort (*i.e*., 20 min/day for five days) and returned twice to test the long-term effects of learning and stimulation. We placed the anode electrode over the right motor cortex (M1), centered over the representation of the first dorsal interosseous (FDI) muscle of the left hand on the motor cortex, identified using single-pulse TMS.

*Figure 4a* shows the main results of this experiment. We tested the placebo groups in the same way as we did the unstimulated cohort in the first experiment and found similar relative differences between age groups. All statistical tests for both experiments are detailed in the *Supplementary Materials*. The first row of *Figure 4b* shows the same lack of offline learning in middle-aged and older participants that was replaced by offline worsening, as seen in Experiment 1 (*Figure 1b*).

**Figure 4.**
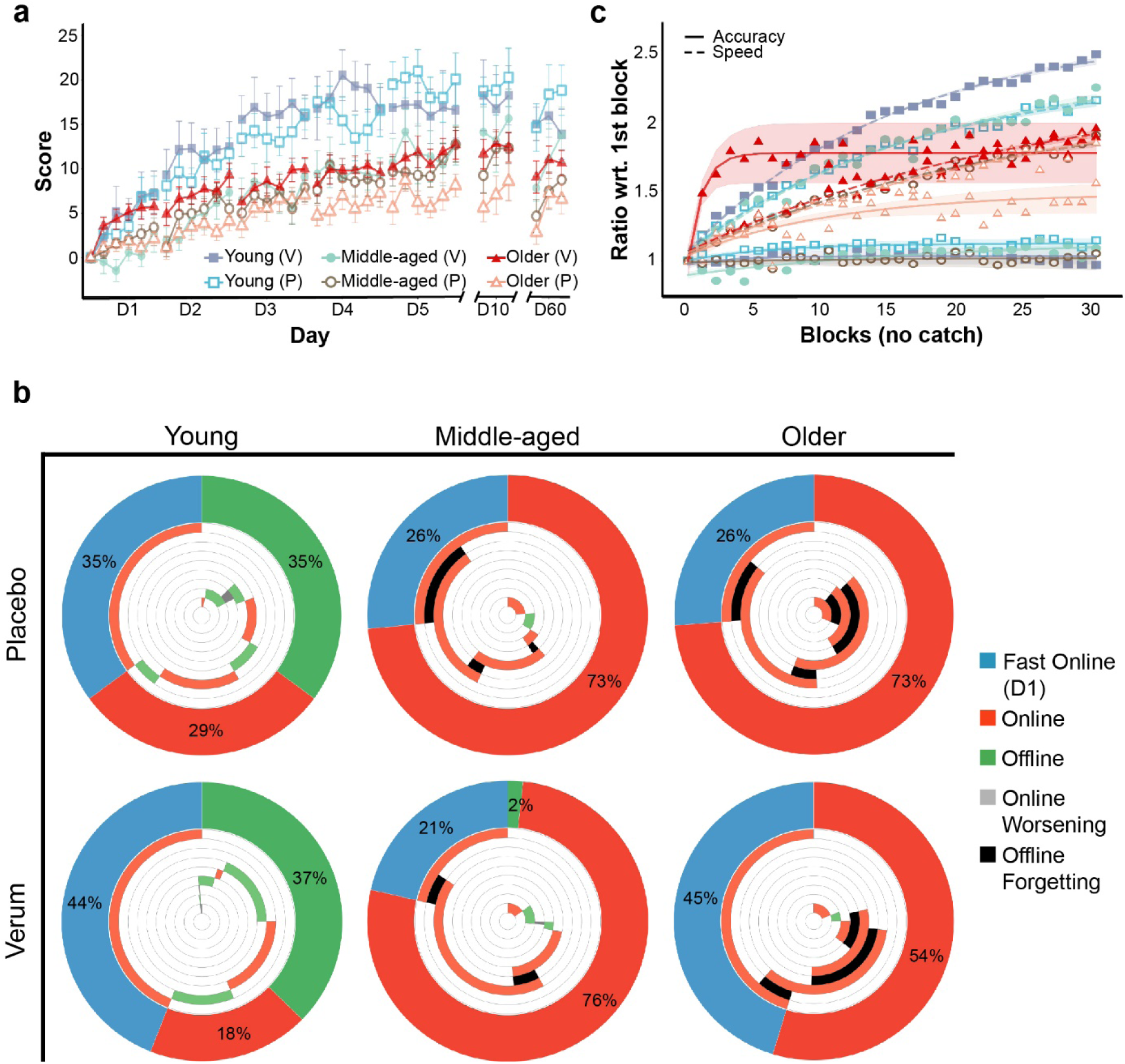
Motor skill acquisition in Experiment 2. **a)** Average scores generated during Experiment 2, consisting of motor training with stimulation, with the error bars depicting the standard error of the mean. The data are grouped by age group and stimulation type (i.e., verum (V) or placebo (P)). **b)** Percentage of total learning over the entire training week represented by fast online learning (D1), online learning during D2-5, and offline learning between training days. Each outer ring captures the proportion of total learning by these three aspects, while the inner rings present their time course during the week (anticlockwise): 1^st^ inner circle is the online performance gain during D1, 2^nd^ inner circle is the offline performance gain between D1 and D2, 3^rd^ inner circle is the online gain during D2, etc. Orange and green represent improvements, while black and gray represent worsening of performance. Please note the large difference in regard to the proportion of total learning explained by fast online learning between the verum and placebo groups in older participants. **c)** Speed and accuracy normalized to the first block of training, grouped by age group and stimulation type (i.e., verum (V) or placebo (P)). Please note that while most groups show similar dynamics to those seen in the respective age-matched groups from the first experiment, the older group receiving verum stimulation shows dynamics more similar to those seen in young adults. As in the first experiment, the accuracy was maintained even at increasing speeds, although young adults receiving verum stimulation significantly dropped in accuracy on the last training day. The shading represents the 95% confidence interval for the logarithmic curve fitting (the type of curve is for display purposes only and not included in the LME analysis.

After verifying the findings from the first cohort, we tested the effects of verum and placebo stimulation in each separate age group. Total learning was not significantly different in the young (T_[20]_ = 2.68, d = 0.53, p = 0.37) or middle-aged (T_[20]_ = 1.58, d = 0.44, p = 0.44) groups. In the older group, however, we found higher total learning in the verum group (T_[24]_ = 4.56, d = 1.53, p = 0.01) with respect to placebo. In the follow-up sessions, we found no significant differences in scores between the verum and placebo stimulation in young and middle-aged participants. Performance on day 60 did not change in the young group with respect to day 5, but dropped significantly in the middle-aged (T_[150]_ = 2.41, d = 0.61, p = 0.003) and older groups (T_[150]_ = 1.4, d = 0.43, p = 0.02). However, the older group undergoing verum stimulation continued to score significantly higher than the group undergoing placebo stimulation (T_[21]_ = 4.92, d = 1.49, p = 0.02). After reconsolidation on day 60 (*i.e.,* after the first block), scores were not significantly different from scores on day 5, providing no evidence for skill loss, but a maintenance of the acquired skill even two months after training.

We did not find a significant effect of stimulation in either online or offline learning when testing all training days. When testing fast online learning as a separate component of learning ^28^, we identified a steeper improvement rate in the older group receiving verum stimulation relative to placebo (T_[21]_ = 3.5, d = 1.15, p = 0.01). We illustrate this difference (and lack thereof in the other groups) in the second row of *Figure 4b,* showing the proportion of the different components of learning to total learning in the verum group (outer rings). These proportions were similar to those in the placebo group for both young and middle-aged adults. In the older group, the proportion of fast online learning to total learning was much larger than that in the placebo group.

*Figure 4c* shows the dynamics of speed and accuracy in the second experiment. There were no significant differences in speed between the verum and placebo groups in any of the three age groups. In terms of accuracy, the young group receiving verum stimulation was significantly less accurate than the placebo group by the end of training (T_[23]_ = 0.07, d = 0.97, p = 0.04). The middle-aged group receiving verum stimulation was significantly less accurate than the placebo group on the first (T_[21]_ = 0.14, d = 1.67, p = 0.002) and second (T_[21]_ = 0.1, d = 1.21, p = 0.02) days. In the older group, the verum group was consistently and significantly more accurate than the placebo group on all days, except the third (please refer to *Supplementary Materials* to find the results of the comparisons), even though we did not find a significant difference in accuracy on the first block (F_[1]_ = 0.38, p = 0.84). As in the first experiment, the older group receiving placebo stimulation reached its maximum accuracy gradually over the course of the week. In contrast, the older group receiving verum stimulation displayed a sharp increase on the first day and quickly reached its plateau, with dynamics reminiscent of those observed in the young group in the first experiment (please see *Figure 1c*, solid lines). In summary, verum atDCS led older participants to score higher by the end of training. Yet, it appears not to have had any effect on the speed of execution of the task; our results suggest that verum atDCS led older adults to improve their accuracy quickly on the first day, much like young adults did in the first experiment.

### Chunking and stimulation

In the first part of the analysis, we proposed that differences in accuracy could derive, at least in part, from differences in the consolidation and deployment of motor chunks during training between age groups. We applied the same classifier trained with data from the unstimulated cohort to the chunking patterns extracted from participants receiving verum and placebo stimulation (data were not seen before by the SVC). *Figure 5a* shows that on the first day of training, the model classified most young participants correctly, while it classified most middle-aged participants as young; this matched the results obtained from the unstimulated cohort. The model also correctly classified older participants receiving placebo stimulation. In contrast, the model classified almost half of the older participants receiving verum stimulation as young. By the end of training, most participants executed chunking patterns more similar to those of the young, matching our previous findings. *Figure 5b* shows the gradual evolution of chunking patterns in all groups, with young adults executing patterns in a consistent manner. Middle-aged adults start with patterns more similar to those of young adults, which become even more similar over the course of the training week. Older adults drift from “old-like” patterns to “young-like” patterns, as seen in the first experiment, with this transition occurring sooner in the verum group.

**Figure 5.**
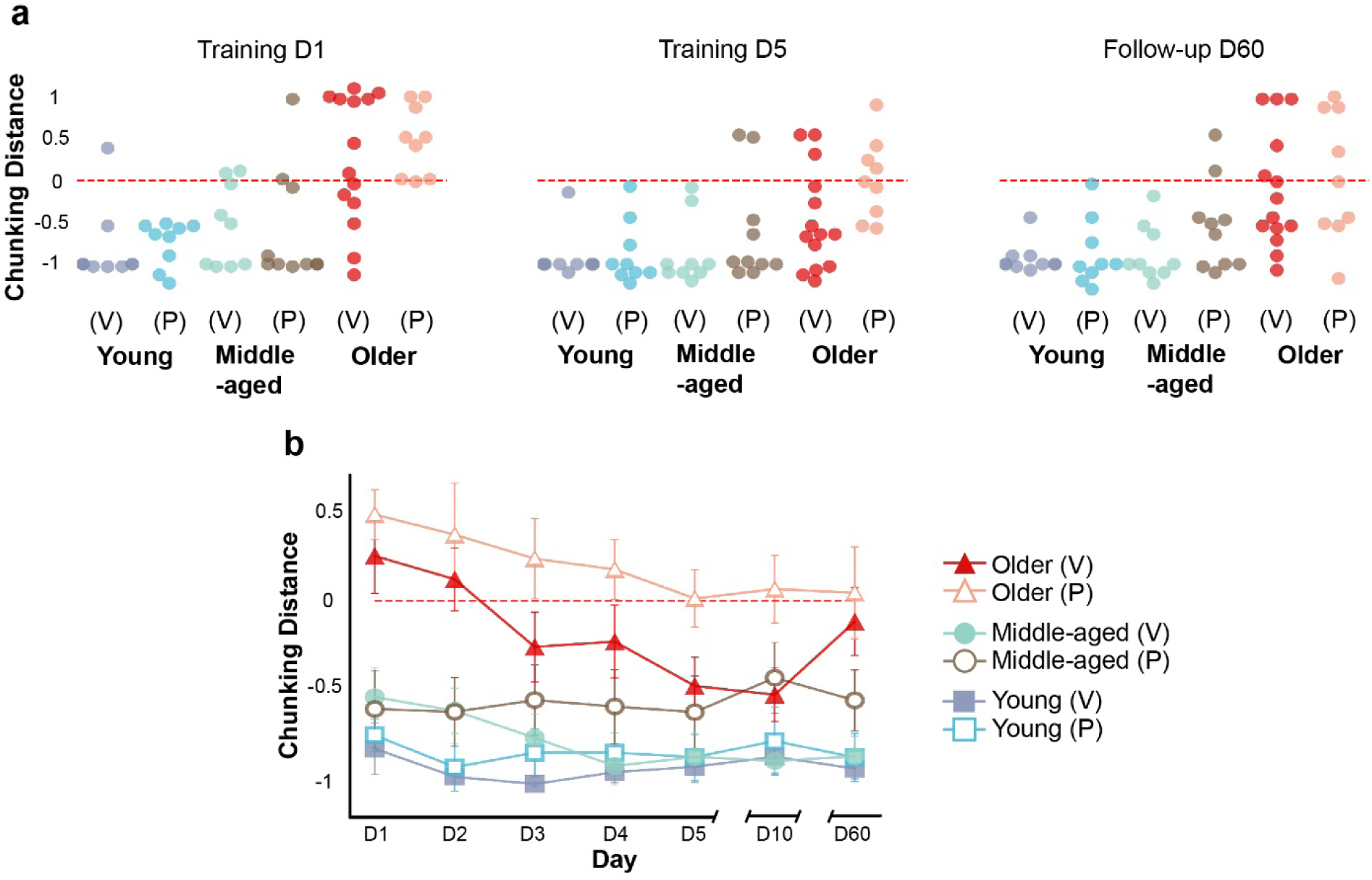
Evolution of chunking patterns during motor skill acquisition in Experiment 2. **a)** Chunking distance for patterns generated on the first and last days of training, as well as on the last follow-up session (i.e., day 60), for each group receiving either verum (V) or placebo (P) stimulation. As described in Figure 3, the red dashed line indicates the boundary between the old (distance > 0) and young (distance < 0) classes. Please note that all groups present a similar set of distances as in the first experiment (Figure 3a), with the exception of the older group receiving verum stimulation, in which half of the participants generated chunking patterns similar to those seen in young adults on the first day of training. **b)** Average chunking distance for each group on each day, with its corresponding standard error. Please note the earlier appearance of “young-like” chunking patterns in the older group receiving verum stimulation compared to the placebo group.

To test whether these differences in chunking translate into different performances in the task for the older group, we identified the participants in the verum group classified as young on the first day and plotted their scores separately from the other participants in their group, as well as the speed and the accuracy. *Figure 6a* shows the scores of all older participants, with “young-like” participants in the verum group scoring significantly higher than those in the “old-like” participants in the verum group (T_[27]_ = 9.36, d = 3.27, p < 0.0001). Older adults generating young-like patterns on the first day of training were significantly faster on the first training block (T_[12]_ = 4.25, d = 1.41, p = 0.02), and overall on the first training day (although this difference was only a trend, (T_[12]_ = 2.77, d = 1.71, p = 0.07)), compared to their peers under the verum condition, but not chunking like young on the first day of training. From the second day onwards, young-like older adults were consistently and significantly faster than their peers. The rate of improvement in speed (i.e., the slope) was significantly steeper in older adults generating young-like chunking patterns on the first day compared to those who did not (T_[80]_ = 0.54, d = 0.29, p = 0.005). As for the accuracy, even after the sub-division based on generated chunking patterns, all older adults receiving verum stimulation improved theirs sharply on the first training day, while older adults receiving placebo stimulation did so gradually over training. Older adults not chunking like young on the first training day (both in the verum and the placebo groups) required more extensive practice to generate young-like chunking patterns, an achievement they reached at different time points of training depending on their speed (please refer to *Figure 6c* and *Figure 6d*), supporting the notion of “a tendency to chunk facilitating rapid execution, and the need for rapid execution inducing chunking” ^40^.

**Figure 6.**
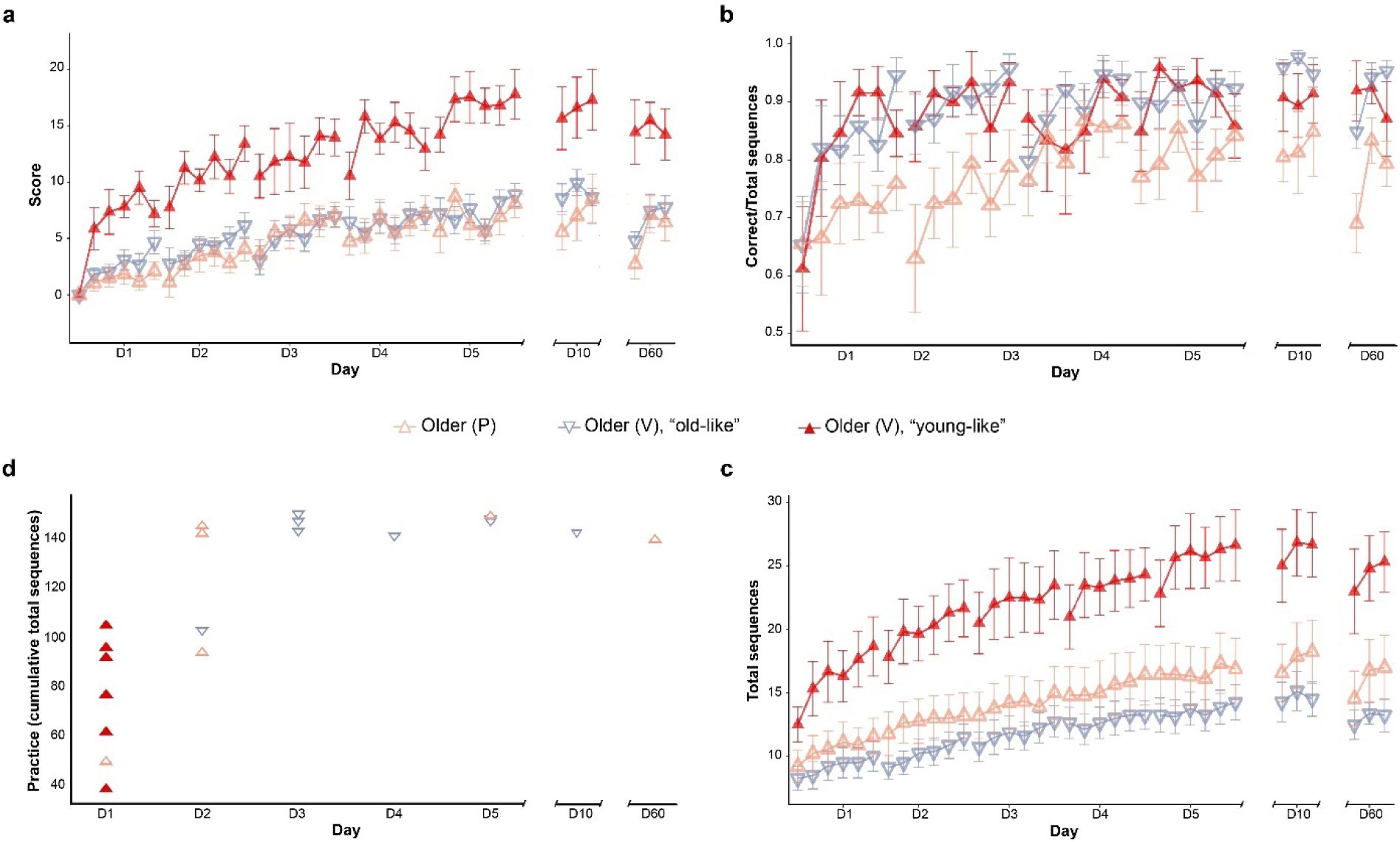
Primary outcome of Experiment 2 for the older groups only, with the group undergoing verum stimulation stratified based on the chunking patterns generated on the first training day. **a)** Older participants in the verum (V) group generating “young-like” patterns on the first day show an enhanced performance compared to those generating “old-like” patterns in the verum group and those in the placebo (P) group. The error bars depict the standard error for scores averaged over each training block. Please note the steep increase in the older group who are chunking like the young group, that is, the increase in performance is reminiscent of the increase seen in participants in the young group. b) Older adults in the (P) and *(V)* groups (for the latter, regardless of whether they generated young-like chunking patterns on the first day or not) started training from comparable levels of accuracy. While the (P) group improved their accuracy gradually over the course of training, young-like and old-like members of the (V) group improved their accuracy sharply on the first training session, at a rate comparable to that seen in young adults of the first experiment. c) Older adults in the (V) group generating young-like chunking patterns on the first day were initially faster than their peers, and increased their speed at a steeper rate. d) This graph depicts the amount of practice, calculated as sum of total sequences generated, up until each individual generated young-like chunking patterns, and on which day they did so. Young-like older adults in the verum group required fewer practice than their peers, most of which required similar amounts of practice (∼140 sequences) to generate young-like patters. Please note that, when considered alongside the speed of each group (i.e., Figure 6c), this suggests that at lower speeds (i.e., those seen in the old-like verum and placebo groups), the required amount of practice is higher than it is at higher speeds.

### Chunking, stimulation, and neurophysiology

We used established TMS protocols to measure intracortical inhibition in all participants in the second experiment. We applied a well-established double pulse TMS paradigm (*i.e.,* short-interval intracortical inhibition, SICI) ^21,41,42^ before and after the first training session to quantify interneuronal GABAergic inhibition within M1, which is directly involved in the learning and execution of the motor sequence. We applied the SICI paradigm while participants were at rest. Within the placebo groups, we found no significant differences in inhibition before or after the first training session or after the whole training week. Similarly, we did not find significant differences between verum and placebo in any of the age groups, thus confirming previous reports that SICI does not significantly change when atDCS is applied together with motor training ^43^.

## Discussion

Here, we studied age-related differences in the acquisition of a sequence-tapping task and applied atDCS concomitant to motor training seeking to enhance performance. We isolated different components of motor skill acquisition intrinsic to this task, and followed their evolution throughout and up to 2 months after training. Applying a black-box testing approach, we contrasted the dynamics of motor skill acquisition seen in young adults, assumed to be able to acquire the task optimally, to those seen in older adults, depicting a generally lower performance on this task. Our results suggest that mastering this motor task relies on the early internalization of the motor sequence, followed by the practice-dependent optimization of its execution, observed as motor chunks at a behavioral level.

The results of the first experiment show that general performance of the sequence-tapping task decreases with age, with score differences between age groups coming mostly from the improvement dynamics present in the first training day. Our results show that speed decreases with age, with relative differences between age groups as expected from natural muscular deterioration ^44^ and atrophy in cortical regions and the corpus callosum ^19^ that occur during healthy aging. Accuracy, on the other hand, was initially comparable among the three groups, but improved sharply and reached a plateau on the first training day in young adults, while older adults improved theirs gradually over the course of training. These results suggest that young adults improve their performance of this task first by minimizing the error of execution and focusing on improving speed thereafter, a behavior reminiscent of the model proposed by Hikosaka and colleagues ^36^.

Instructing individuals to generate sequential movements in quick succession results in the spontaneous appearance of temporal patterns known as motor chunks ^6,7^. Our analyses on the chunking patterns produced by all participants showed that older adults did not generate chunking patterns as young adults did on the first training day, but did so after more extensive practice. Previous research on motor chunking assumes chunks emerge from the repeated sequential execution of single commands in close temporal proximity ^9^. In our first experiment, young adults were faster, so one could deduce that this higher speed allowed for more intensive practice on the first day of training and that this led them to generate chunking patterns sooner.

The second experiment revealed a significant effect of atDCS only in older adults, with those under the verum condition scoring significantly higher than their peers in the control group. The same analysis performed on the data from the first experiment revealed that older adults receiving verum atDCS during training reduced their error sharply on the first training day, much like young adults of the first experiment did. Additionally, they generated chunking patterns similar to those seen in young adults at earlier stages of training, with many of them doing so on the first training day. At a first glance, these results would suggest not all older adults responded to atDCS to the same extent. However, the speed and the accuracy of older adults under the verum condition, grouped according to whether they generated young-like chunking patterns or not on the first day of training, suggest that atDCS acted on all older adults by facilitating the encoding and the storage of the sequence in memory, leading to a sharp improvement in accuracy on the first training session. On the other hand, older adults with young-like chunking patterns were faster initially, which alongside an optimized accuracy achieved on the first training day, resulted in an earlier consolidation of motor chunks.

Therefore, older adults under the verum condition improved their performance following a similar pattern as the one seen in young adults, optimizing the error first and improving the execution thereafter, albeit at different rates depending on their speed of execution. As previously mentioned, speeding up leads to chunk formation, and chunk formation allows further increases in speed, which is supported by faster, young-like older adults chunking earlier and increasing their speed more steeply.

### Motor and cognitive components of the sequence-tapping task

Recent work has suggested that motor chunks are not represented within M1, but rather form in premotor cortical and striatal centers ^45^, with patterns represented in the parietal cortex, as well as in dorsal and ventral premotor cortices ^13^. Subsequent chunk selection occurs in the striatum ^45^ and bilateral putamen ^46^, as well as dorsal premotor and supplementary motor areas ^47,48^, with chunk execution eventually occurring in M1 ^14^. Krakauer and colleagues ^15^ suggest that the lack of representation of motor chunks within M1 indicates these structures are not selectively motor, but rather cognitive elements independent of motor execution, whose function is limited to storing the order of the sequential elements for their efficient retrieval. If this were the case, it would suffice to memorize the sequence for such patterns to emerge and, in the case of the present study, to optimize its execution and minimize the error. In the real world, this would translate to a pianist being able to master a musical piece simply by studying the score, which appears not to be the case ^49^.

Our method captured behavioral differences in speed, accuracy, and generated chunking patterns. Considering speed increases consistently in all groups without accuracy ever decreasing, the observed improvement results from a shift in the speed-accuracy trade-off ^4,50^. The increase in accuracy likely reflects the storage of the sequence elements in memory, resulting from the transition from a state of high uncertainty (i.e., ignorance of the sequence) to a state of low uncertainty (i.e., knowing the sequence). This information constitutes the spatial feature set, specific to the trained sequence ^48^, and likely enables an increase in speed (without sacrificing the accuracy) by boosting motivation and confidence in the execution itself ^51^. This process would indeed be independent of motor practice, and would capture the cognitive dimension of this task.

The chunking patterns, as detected by our method, reflect a different set of features. As the present method uses the inter-key intervals to identify the chunks, it portrays the rhythm of execution, which constitutes the temporal feature set. Specific temporal patterns can be encouraged externally (for an example, refer to this experimental paradigm ^52^), much as the partition determines the tempo in music playing. Taught patterns can be suboptimal, requiring the execution of relatively difficult transitions in close temporal proximity, but in the absence of external temporal cues, easier transitions are normally grouped together ^53^. These patterns are optimized with practice, and their structure is constrained by a balance between computational cost and motor efficiency ^38^.

Kornysheva and Diedrichsen ^52^ found neural activity encoding spatial and temporal features independently represented in lateral and bilateral medial premotor cortices. These findings suggest the emergence of motor chunks, as those captured by our method, result from the storage of both spatial and temporal features in higher brain areas, upstream from M1 in the motor network, a mechanism that appears to be diminished in older adults. Our results suggest that the most effective way to improve performance in this task is to first store the spatial components (i.e., sequence order), followed by the storage and iterative optimization of the temporal features (i.e., chunking patterns). This order may be specific to the present task, in the sense that no external temporal cues were provided; indeed, it may be that when both the sequence order and the temporal patterns are explicitly available, both may be consolidated in parallel.

### Black-box testing

There is an ongoing discussion on whether motor chunks are purely cognitive elements ^15^. In the presently discussed analysis, we consider young adults to be capable of acquiring motor skills optimally (within the constraints inherent to the human neuromotor system). As such, we consider them to embody the requirement of the system for the acquisition of our sequence-tapping task. In young adults, the accuracy reaches a plateau in the early stages of training, indicating the sequence has been stored in memory. Chunking patterns, on the other hand, also emerge on the first training day and remain relatively unchanged for the rest of training. This early optimization of both the accuracy and the chunking patterns could be interpreted as chunking patterns being a direct consequence of storing the sequence in memory. However, the process we observe in older adults suggests otherwise. In older adults, the mechanism for storing the sequence in memory appears to be diminished, as their accuracy increases gradually over the course of the training week. Nevertheless, atDCS seems to restore this mechanism early in training, similar to reactivating a dormant ability of the brain, leading to the early consolidation of spatial features and resulting in an optimized accuracy on the first training session, as seen in young adults. Chunking patterns appeared at different stages of training, which seems to depend on the amount of practice and, indirectly, on the speed of execution. *Figure 6a* shows faster older adults chunking sooner, which would support the notion of increased amounts of practice early in training leading to an earlier consolidation of chunks. This is not supported by the other older adults, as most of them need similar amounts of practice to generate young-like chunking patterns, regardless of differences in speed; please note that patterns emerging later in training, but requiring similar amounts of practice imply a slower execution. It appears that relatively high speeds (e.g., like those seen in young and middle-aged adults) place a prime on the optimal execution of the sequence, resulting in an accelerated formation of chunking patterns, while executing the sequence at lower speeds (e.g., old-like older adults under verum and all older adults under the placebo) leads to a slower consolidation of chunks, requiring more extensive practice.

These results support the notion of a critically-important cognitive component intrinsic to sequence-tapping tasks ^15^. However, the structure of the patterns detected as motor chunks at a behavioral level is determined by the ease of the mechanical transition between key presses ^53^, and is optimized with practice (a process occurring more gradually in slower older adults). This might explain why pianists need extensive practice to perfect a musical piece, long after memorizing it.

### The effect of atDCS on the storage of spatial coordinates

Our results suggest atDCS facilitates the consolidation of motor chunks in older adults when the anode is placed over the contralateral hand-knob representation at M1 (though it is of note that the spatial resolution of the stimulation is limited) by facilitating the storage of the sequence order in memory. Given the non-focality attributed to this technique ^54^, we cannot discard the possibility of physiologically relevant stimulation of other brain areas, like the premotor cortex, as spatial and temporal components of the sequence are encoded unilaterally and bilaterally, respectively, in these regions ^52^. Therefore, the anode could be inducing LTP-like plasticity ^55^ in intracortical interneurons ^56^ of the M1 contralateral to the trained hand, but also the ventral and dorsal premotor areas (i.e., PMv and PMd), facilitating the storage of the spatial coordinates in these regions.

In search for likely causes behind the selective effect of atDCS, seemingly exclusive to older adults, we quantified intracortical (GABAergic) inhibition within M1, since previous studies have shown that less efficient SICI was associated with lower dexterity in executing rapidly alternating two-finger tapping ^20,21^. Nevertheless, we did not find significant differences between verum and placebo in any of the age groups, confirming previous reports that SICI does not significantly change when atDCS is applied together with motor training ^43^. This further supports the view that the effect of atDCS in older adults at a behavioral level does not directly act on the execution of the sequence itself, but likely on higher brain areas, upstream of the motor network.

If we conceive the process of chunk formation as the transition from high-uncertainty to lower-uncertainty in the execution of a motor sequence, we can consider each source of information to contribute to this change of state. In a recent study, Cross and colleagues found evidence for the primary motor cortex being a hub where both somatosensory and visual feedback converge ^57^. This information is likely integrated in higher brain areas, such as premotor ^58^ and parietal cortices ^16^, an essential process to decrease the error in the execution of our task. Sensorimotor integration decreases with age ^59^, so atDCS could be compensating this process in older adults.

### Implications for using atDCS in motor sequence learning

During the first day of training, young adults optimize their accuracy and focus on improving speed thereafter, experiencing a shift in the speed-accuracy tradeoff early in training. The consolidation of spatial coordinates allows the rapid optimization of temporal features, resulting in an early consolidation of chunking patterns. The fact that these patterns do not change much during the subsequent four days of training indicates that they reached an optimized strategy. As such, these strategies would constitute a ceiling on dexterous skill for this task. Our results suggest that atDCS influences accuracy, but not speed, as we did not find significant differences in speed related to stimulation in any age group. Nevertheless, it appears that only imbalanced neural systems that are less than optimal can benefit from stimulation. Neural responsiveness decreases with healthy aging, which is why enhanced plasticity induced by atDCS ^60^ likely benefits older adults ^3^. Greeley and colleagues ^61^ reported improved motor chunk formation related to atDCS applied to M1 in young adults. Unfortunately, we cannot compare our results to theirs, as the metric they use is the number of chunks, which does not provide much information on how the sequences are segmented. Further, they consider fewer chunks to reflect greater improvement, which is based on the notion of an eventual full concatenation of chunks, which likely is not attained due to the related computational costs ^38^. On the other hand, they did not find differences in speed nor accuracy, which matches our own findings.

## Limitations

atDCS applied to enhance motor performance has low focality ^54^, which limits the interpretation of how exactly stimulation restores motor and cognitive functions in older adults in regard to which brain areas of the motor network are mainly involved. Additionally, the fact that the electrophysiological evaluation (i.e., TMS) could not be achieved in all subjects limits the null-finding of SICI in relation to learning and atDCS in this study. In regard to the statistical analysis, we understand there are certain implications to centering the data, but as we use these comparisons just to lay down the grounds for the discussion on the actual analysis on the mechanisms behind apparent differences in motor skill acquisition, we consider the correction and corresponding statistical tests are justified.

## Conclusions

Sequence learning is essential and ever present in the execution of many activities of daily life. The black-box approach, contrasting different components of skill acquisition between young and older adults, whose ability to acquire new skills is present but diminished, has revealed that mastering motor sequence tasks depends on the early storage of the sequence elements in memory (i.e., consolidation of spatial components), leading to the practice-dependent emergence of temporal patterns (i.e., motor chunks). Non-invasive brain stimulation, as applied here, might support and accelerate this process in systems not working at the optimal level, such in healthy older adults.

## Methods

### Participants

A total of 113 subjects volunteered to participate in our study, categorized as young (18-30 y/o; n = 41, 27 female; age*μ* = 24.5 y/o), middle-aged (50-65 y/o; n = 34, 20 female; age*μ* = 57.7 y/o), and older (>65 y/o; n = 38, 21 female; age*μ* = 72.3 y/o) healthy participants. All participants were right handed as determined by the Edinburgh Handedness Inventory ^62^. The participants reported not having a previous history of serious medical conditions (General Health Questionnaire, GHQ) or contraindications for tDCS and TMS (questionnaire based on safety recommendations for these techniques ^63^). We performed a neurological examination on all participants over the age of 50 to ensure that participants were healthy and performed the Mini-Mental State Exam (MMSE, ^64^) to ensure that all participants scored at least 26 out of 30 points. The participants gave their informed consent under protocol guidelines approved by the cantonal ethics committee Vaud, Switzerland (project No. 2017-00301) and the ethics committee Hamburg, Germany (PV 3770) according to the Declaration of Helsinki.

### Motor task

We used a well-established finger-tapping task ^2,3^ that required the participants to replicate a nine-digit numerical sequence displayed on a screen as quickly and as accurately as possible, using a four-button box with buttons labeled from “2” to “5” (“2” for the index finger, “5” for the pinky finger). A white dot on the screen, displayed beneath the numbers, indicated the button to be pressed next. The dot would move to the next digit as soon as a key was pressed, regardless of whether or not it was pressed correctly. Before starting the first training session, we asked all participants to perform a 90-second familiarization block to use as a reference for general initial skill level (please refer to *Supplementary Materials* for details on how we used this sequence). Training started immediately after this block. The participants trained their left hand for 20 min each day for five consecutive days. Each day of training consisted of seven 90-second blocks interspaced by 90-second rest periods. Six of the blocks from each day contained the same sequence (*i.e.,* training sequence). The seventh block consisted of a “catch” block presented halfway through training on each day and contained a sequence different from the training sequence. Each day of training had a catch block with a different sequence. We used the catch blocks to test whether the observed improvements were specific to the training sequence or generalizable to any sequence. We presented the catch blocks at different stages of the training session on each day, alternating between the third, fourth and fifth blocks to avoid interfering with overnight consolidation of learning and anticipation of its appearance ^2^. The participants returned for follow-up sessions on the 10th and 60th days after the beginning of training, during which they executed three blocks of the training sequence; we used these visits to test for long-term retention related to the intervention. We did not provide any form of feedback on the participants’ performance at any time.

### Electrophysiological exploration of changes within M1

We used TMS to identify the representation of the FDI muscle of the left hand and quantified the interneuronal GABA_A_ receptor-mediated inhibitory networks within the right M1. TMS was delivered with a 70-mm figure-of-eight coil linked to a Magstim BiStim2/Magstim Bistim machine (Magstim Ltd., Whitland, UK), and we recorded electromyography (EMG) signals from the FDI muscle. We first empirically identified the cortical target as the spot on the scalp eliciting the largest motor evoked potential (MEP) under EMG control in the left FDI. Then, we identified the resting motor threshold (RMT) as the minimum single pulse intensity to evoke 50-µV MEPs 50% of the time and the single pulse intensity to evoke 1-mV MEPs (test intensity). The SICI was quantified with a well-established paired-pulse paradigm using a conditioning pulse delivered at 80% RMT intensity followed 3 ms later by a test pulse delivered at the test intensity ^21,41,42,65–67^. We assessed the RMT, 1-mV test intensity, and SICI before and after the first training session, after the 5th day of training, and on the 60th day after long-term retention testing.

### Electrical stimulation of M1 during training

When applying atDCS, the anode was placed over the FDI hotspot, and the cathode was placed over the left supraorbital area ^3^ using squared electrodes (25 cm^2^), covered in sponges soaked in saline solution (0.9% NaCl), connected to a neuroConn DC-STIMULATOR (Germany recordings) or a DC-STIMULATOR PLUS (Switzerland recordings) (neuroConn GmbH, Ilmenau, Germany). Stimulation was applied in a double-blind, placebo-controlled, parallel design, with all experimenters involved in the acquisition and/or the analyses of results blinded until the end of the acquisition. The verum stimulation consisted of 20 min of stimulation with 1 mA direct current (ramp-up/down times of 8 seconds). The placebo stimulation consisted of 40 seconds of stimulation delivered at the beginning of training (with 8-s ramp-up and 5-s ramp-down times, as defined by neuroConn) to emulate the prickling sensation on the scalp often reported in the use of this technique during current intensity variation ^68^.

### Experimental protocol (*Figure 7*)

The participants in both Experiment 1 and Experiment 2 came for seven visits. The participants in Experiment 1 started with motor training, while the participants in Experiment 2 started the first day with a set of electrophysiological investigations: identification and neuro-navigated registration of the FDI hotspot coordinates, identification of the RMT and 1-mV intensity, and a battery of 24 single and 24 double TMS pulses. After these measurements, the participants executed the motor training described above with concomitant atDCS for 20 min. We repeated the session of electrophysiological investigations after the first and fifth training sessions and the 60^th^ day control session. For each of these investigations, we adjusted the intensity of the test TMS pulse to maintain a 1-mV amplitude of the single-pulse MEP.

**Figure 7.**
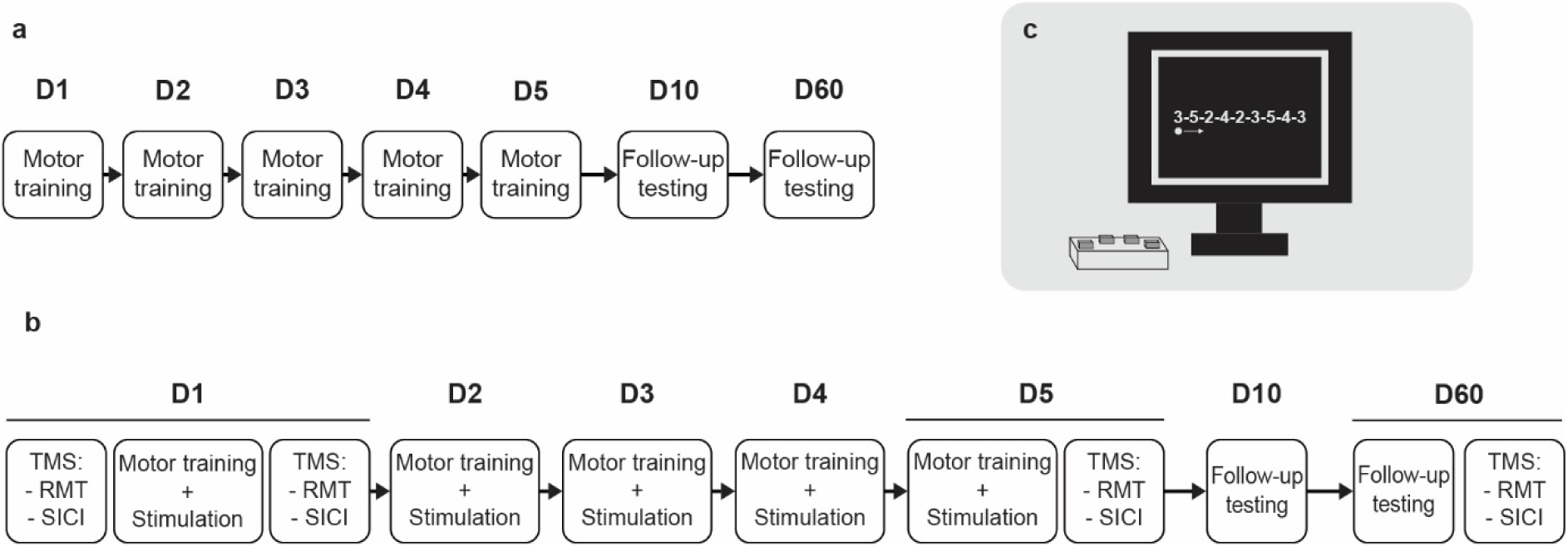
Experimental protocol. a) Experiment 1 tested the behavioral outcome of five days of training in three groups: young, middle-aged, and older healthy adults. b) Experiment 2 tested the behavioral and electrophysiological outcomes of five days of training with atDCS delivered to the motor cortex in six different groups: young, middle-aged, and older healthy adults receiving either verum or placebo stimulation in a double-blind parallel design. c) The motor training consisted of pressing, as quickly and accurately as possible, four buttons corresponding to the non-opposable fingers of the left hand (5 = pinky finger) according to an explicit sequence displayed on a computer screen.

### Chunking strategy extraction

We extracted a single chunking pattern to characterize the execution of the training sequence for each day of every participant’s training. To this end, we applied the clustering approach proposed by Song and Cohen ^69^ and labeled successive interkey intervals (IKIs) of every sequence as either “fast” (*i.e.*, “1”) or “slow” (*i.e.*, “0”) considering adjacent keys with intervals labeled “fast” to belong to the same chunk. Please refer to the *Supplementary Materials* for a detailed exposition of our arguments in favor of using this approach. Each sequence had nine IKIs, with the first reflecting the interval between the last key press of the previous sequence and the first key press of the current sequence. After removing incorrect sequences from each block, we normalized the IKIs in each sequence to the total duration of the sequence (*i.e.,* divided each IKI by its sequence duration) to account for the gradual increase in speed during training. After normalization, we applied the K-means clustering algorithm (Sklearn, https://scikit-learn.org/) to sequences of each block, enforcing the notion of two clusters being present (*i.e.*, “fast” and “slow”) by labeling the IKIs in each sequence based on their proximity to them. The outcome of this step was a chunking pattern for each individual sequence. To determine a single pattern describing strategies for each day, we defined a series of possible criteria:

1. Maximum allocation: This criterion looks at the most frequently repeated chunk sizes generated by a participant and excludes patterns with chunk sizes different from these. It also assumes that participants will allocate all keystrokes to at least one of the chunks. These two constraints result in the choice of one dominant pattern for each day.
2. Reclustering: This criterion sums all chunking patterns for each day and reclusters them using K-means clustering, outputting a single pattern for the entire day.
3. Reclustering top: Similar to the previously mentioned clustering approach, with the difference that it reclusters only a percentage of the most frequently repeated patterns. In this case, we fixed this percentage to 15%.
4. “More-often-than-not”: This criterion uses all chunking patterns generated on a day and generates a new sequence containing “1”s for each IKI labeled “1” in more than 50% of the sequences of that day and zeros otherwise.
5. Highest frequency: This criterion takes the most frequently repeated pattern on each day.

For some participants, the pattern found to characterize their execution for a given day varied depending on the criterion used. For this reason, we used all five criteria and generated a single chunking pattern by performing a majority vote on the five patterns. In other words, we obtained five chunking patterns for every participant on each day. Then, we performed a majority vote for each bit (*i.e.,* each IKI label) of the five chunking patterns and obtained a single pattern characterizing chunking on that day. Please refer to the *Supplementary Materials* for more information on this process.

### Chunking pattern classifier

We fitted a support vector classifier (Sklearn, https://scikit-learn.org/) to chunking patterns generated on the first day by young and older participants of the first experiment (*i.e.*, training without stimulation). We used 80% of these chunking patterns as the training set and the remaining 20% as the test set. We did not keep any of these data as the validation set, as we intended to use the patterns generated by young and older adults receiving placebo stimulation in Experiment 2 (*i.e.*, training with stimulation) as the validation set. To fine-tune the model, we performed a grid search cross-validation on different parameters, namely, the regularization parameter (*C*) and the model kernel, and chose the model yielding the highest F-scores in both cross-validation and testing. We repeated this process ten times, varying the samples used as training and testing datasets. After this step, we obtained ten models with parameters optimized to the training set used each time. Among these ten models, we chose the one with the highest F-scores, with the optimal parameters being C = 0.1 and a linear kernel, for which the training F-score was 0.88 and the test score was 1. We chose this model as the final model and used it to classify chunking patterns generated on the second day onwards in the first experiment and all days in the second experiment. As previously mentioned, we validated this model with the chunking patterns generated by the young and the older groups receiving placebo in Experiment 2, with a classification accuracy of 88.88% (*i.e.,* F-score of 0.8888).

We used the decision boundary from the final model, separating the “young” and “old” classes, to quantify the resemblance of chunking patterns from every individual to each class. Specifically, we obtained the distance from each nine-dimensional data point (corresponding to nine IKIs) to the nine-dimensional hyperplane separating both classes and used this amount to assess changes in chunking strategy during training.

### Statistical analysis

We performed all statistical analyses in R ^70^. We used the *lme4* package from Bates and colleagues ^71^ to fit LME models to our data, and we used the *emmeans* toolbox ^72^ for post hoc testing. For the effect sizes, we used the calculation implemented in *emmeans*, which looks at pairwise differences and divides them by the standard deviation, and used confidence intervals to account for uncertainty in estimated effects and estimated standard deviation. We fitted all models using restricted maximum likelihood (REML). We tested the significance of fixed effects by means of ANOVA Type III on the model using Satterthwaite’s method, and obtained p-values using the *lmerTest* package ^73^. We performed post hoc tests on significant fixed effects and corrected for multiple comparisons using Tukey’s HSD method ^74^. We ran two-tailed post hoc tests on the estimated marginal means (*i.e.,* least-squares means) from our fitted models, with degrees of freedom estimated using the Kenward-Roger method ^75^. The present manuscript discusses, with a few exceptions, significant results only (with a cutoff for statistical significance of p < 0.05). Please refer to the *Supplementary Materials* for the results for all statistical tests applied to the data from both experiments.

## Supporting information

Supplementary information on scoring, statistical tests, uncorrected data, chunking pattern extraction, etc.

## Data Availability

The datasets we acquired during both experiments discussed in this study are available from the corresponding author upon reasonable request.

## Acknowledgment

The project was partially funded by the Defitech Foundation (Morges, CH), the Federal Ministry of Education and Research (BMBF 01GQ1424B, GER), the Wyss Foundation (Geneve, CH), the German Research Foundation (SFB 936-C4) and the Medical Research Fund of the University of Hamburg (GER).

## Author Contributions

Conception (FCH); protocol development (PME, JET, ACS, TM, MJW, FCH); data acquisition (PME, TP, JET, ACS, MJW); data analyses (PME, TP, ACS, TM, MJW, FCH); interpretation of results (PME, TP, JWK, MJW, FCH); manuscript (PME, TP, FCH); manuscript critical revision (PME, TP, JET, ACS, JWK, TM, MJW, FCH); funding (FCH).

