## Supplementary information on scoring, statistical tests, uncorrected data, chunking pattern extraction, etc. for "Black-box testing in motor sequence learning"

### Supplementary Materials

#### Scoring of the behavioral task

We instructed participants to execute the sequence as fast and as accurately as possible, so we tried to score performance in a way that would consider both aspects. Dan and colleagues [1] proposed a measure multiplying the exponential of speed times the exponential of accuracy:

$$PI = \exp\left(-\left(\frac{BlockDuration}{12}\right)\right) * \exp\left(-\left(\frac{Errors}{12}\right)\right) * 100$$

In their equation, the “12” was the maximum number of correct sequences possible in their task per block, and Errors = 12 - CorrectSequences. We tried using this equation. In our case, it was the time that was fixed (*i.e.* BlockDuration) and the total number of sequences per block varied. We noticed that the equation weighted errors less severely if made at higher speeds, while weighting participants making mistakes at lower speeds more harshly. As data came from people from multiple age groups, some of them moving more slowly (*e.g.,* older adults) we chose not to apply this asymmetric weighting. Additionally, we noticed that the time constant used greatly changed the shape of the learning curve (*e.g.,* using either 90 seconds or 1.5 minutes).

We thought about a hypothetical case in which two persons generated five correct sequences, with the difference that the first person generated a total of 10 sequences (50% accuracy) and the second a total of 5 sequences (100% accuracy). Based on what we instructed in terms of speed and accuracy, we wanted to give higher credit to the person having higher accuracy. Directly multiplying speed and accuracy, namely 10*0.5 for the first and 5*1 for the second person, would yield scores of 5 for both. Therefore, we chose to multiply the number of correct sequences times the accuracy, which would yield 5*0.5 = 2.5 for the first and 5*1 = 5 for the second, placing the desired prime on both parameters. The resulting equation is:

$$score = CorrectSequences * PercentCorrect$$

*Correction for individual skill level at the beginning of training*

We used a single baseline block as a benchmark (containing an independent sequence, the same for all subjects) for evaluating the initial motor capabilities of each subject. We do not use this block to normalize the performance during training under stimulation, for several reasons. First, it is a different sequence from the sequence people train on, and even if it is similar in structure and equivalent in complexity, previous research shows there is no transfer between different sequences [2], which we also see when comparing the training blocks to the catch blocks in this study (please see *Table 1*). Second, the training kicks in from the very first execution of a sequence [3]. Third, the performance averaged over the whole block might benefit already from the stimulation, so penalizing it with a different sequence executed under different circumstances (i.e., without stimulation) does not seem adequate.

We used the baseline block as reference for initial skill levels. In the first experiment, we found significant differences in the scores of this block between age groups. Nevertheless, we found no significant differences in accuracy (please see *Table 1*). In the second experiment (placebo groups), we found no significant differences in neither the scores nor the accuracy in the baseline block, further supporting the notion of participants starting at the same skill level (please see *Table 2*).

As we confirmed all participants were healthy, we assumed all age groups could improve within the same range of values, allowing us to compare them within the same frame of reference. For this reason, we corrected individual performance by subtracting the score of the first training block. As participants' improvement is in the same order of magnitude, we consider absolute improvement to be an adequate measure.

### Catch block scores

In the main manuscript, we mention that we see no generalization of learning to sequences different from the training sequence. The scores below show average scores of all blocks, including the catch blocks (please see *Figure 1*).

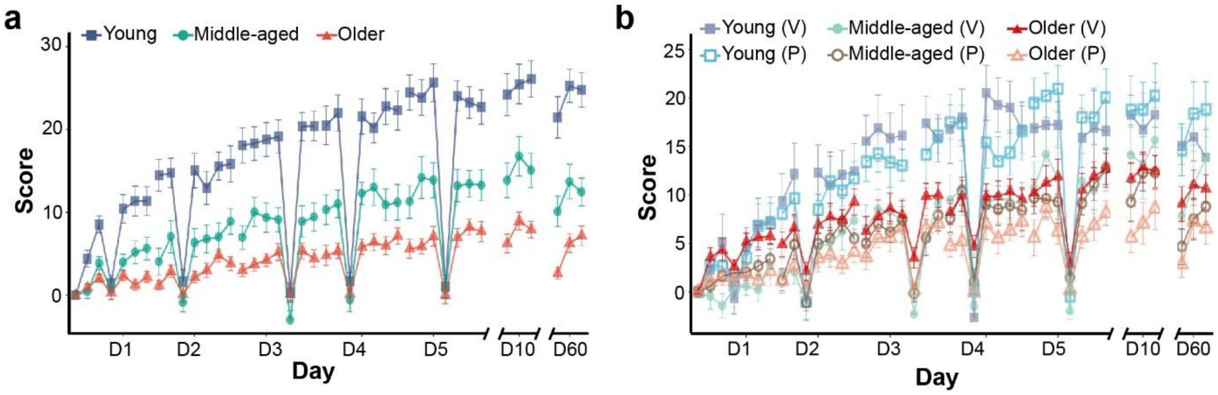


***Figure 1. Primary outcome of all blocks (i.e. including catch blocks). a)*** *Average score for all blocks of the first experiment (i.e. training without stimulation). The vertical lines represent the standard error of the mean.* ***b)*** *Average scores for all blocks in the second experiment (i.e. training with either verum (V) or placebo (P) stimulation), with vertical bars describing the standard error of the mean. The error bars depict the standard error of the mean.*

It is clear to see that the improvement seen in the training sequence does not transfer to other sequences. Scores were not significantly different between catch blocks (*Table 1 and Table 2*).

### Speed and accuracy

In the main article, we only show the normalized speed and accuracy for each group. The reason is that we were interested in discussing the dynamics of both parameters, and intended to show both processes in the same plot. Here, we show the average speed and accuracy for each group. We describe the speed as the total number of sequences generated within a block (*Figure 2*).

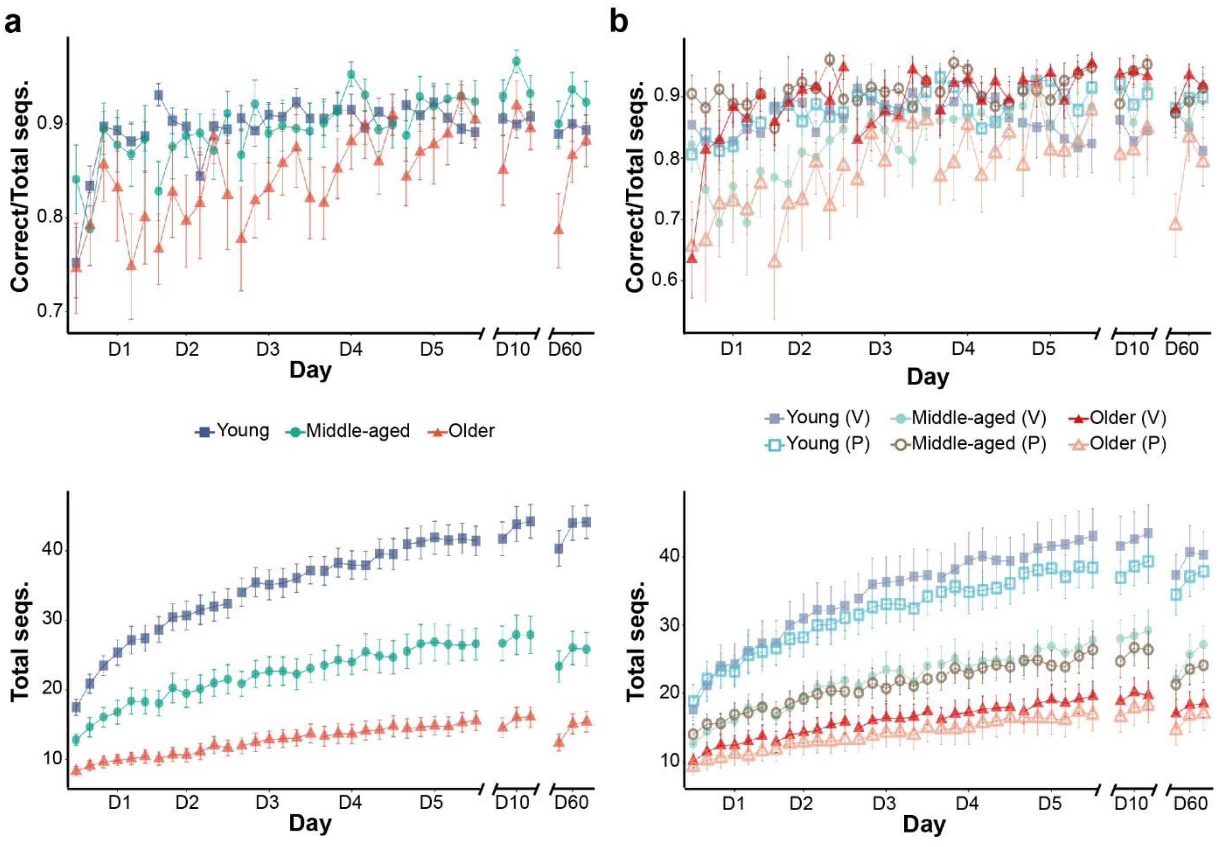


***Figure 2. Group average accuracy and speed.*** *Accuracy (top) and speed (bottom) in the first (a) and second (b) experiments. We defined speeds to be the total number of correct sequences generated within a block, while accuracy corresponds to the ratio of correct sequences to total sequences in each block. The error bars depict the standard error of the mean.*

### Statistical analysis of the main results

At the end of this document, we include tables with all statistical tests we ran. Whenever between-subject variability warranted it, we used linear mixed effect models with random intercept and slopes for individuals. We defined the factor “Day” as categorical, to fit individual lines per training day.

### Uncorrected scores

In the main manuscript, we present centered average scores for all groups of both experiments, which was necessary to compare the scores statistically. *Figure 3* shows the uncentered scores. Interestingly, the unstimulated groups and the groups receiving placebo stimulation look almost exactly the same, and the absence of an effect of atDCS in young and middle-aged adults is more evident.

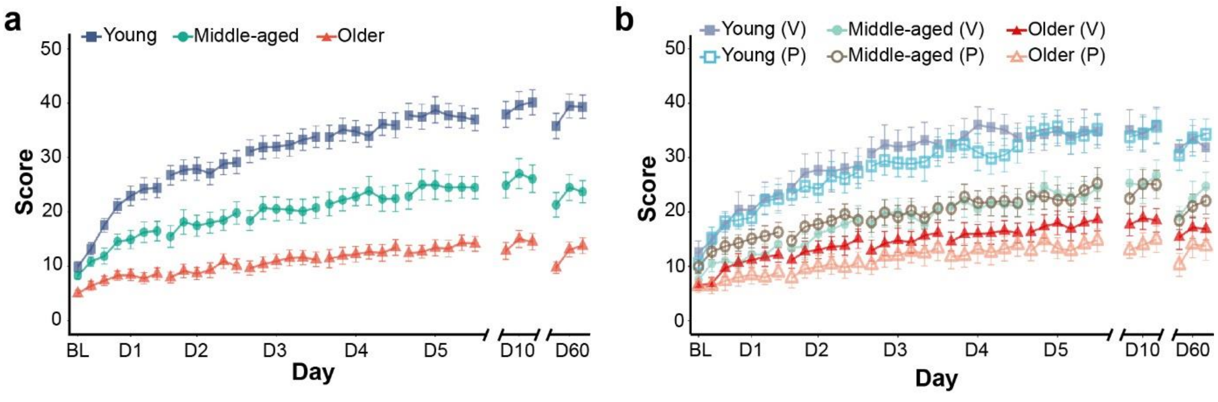


*Figure 3. Uncorrected scores for both experiments. The scores obtained by the three groups of the first experiment (a) match those obtained by the groups receiving placebo stimulation in the second experiment (b). Please notice that without the correction, the verum and placebo groups of both young and middle-aged adults are even more similar, highlighting the absence of an effect reported in the main manuscript. In contrast, older adults receiving verum stimulation outscore the unstimulated and the placebo groups of older adults.*

### Motor chunk estimation from behavioral data

A common approach at extracting chunking patterns in the execution of discrete sequence production tasks is to average the inter-key intervals (IKIs) and detect statistically significant increases, assumed to separate adjacent chunks [4–6]. Acuna and colleagues [7] implemented a probabilistic approach meant to estimate the likelihood of an individual performing a certain chunking pattern based on the IKIs and the errors generated during the execution of the task, as well as the correlation of these two components. The model they propose uses expectation maximization to estimate the most likely chunking pattern generated on each trial (*i.e.* sequence execution). For each trial (t), the model uses prior knowledge on the chunking pattern executed in the preceding trial (t-1). This approach stands on an assumption that makes its use difficult in our dataset. The model assumes that improvement is steady and consistent during training, with similar chunking patterns between adjacent trials. This could be the case in young adults, with average scores monotonically increasing between days. In contrast, middle-aged and older adults often show diminished performance at the beginning of a training session with respect to the previous day. On the other hand, the use of a catch block presented in the middle of training sessions often disrupted performance in the subsequent training block (as reported by several participants and as observed in the “dips” in the learning curves of each day, *Figure 1*), which probably had an impact on the executed chunking pattern. Additionally, errors in older adults, more frequent than in young, would likely obstruct finding the chunking pattern implemented by a participant, as errors cause participants to slow down [8]. Furthermore, the probabilistic approach enforces the notion of chunks eventually being fully concatenated by the end of training, while chunk formation is likely constrained by the computational cost of retrieving a certain amount of sequence elements [9].

For these reasons, we decided to use the approach proposed by Song & Cohen [10]. In their method, chunking strategies are detected by applying a k-means clustering algorithm to the IKIs, forcing two clusters to label IKIs as either “fast” or “slow”.

Each sequence had nine IKIs, with the first one reflecting the interval between the last key press of the previous sequence and the first key press of the current sequence. After removing incorrect sequences from each block, we normalized the IKIs of each sequence to the total duration the sequence (*i.e.* divided each IKI by its sequence duration), to account for the gradual increase in speed during training. After normalization, we applied the K-means clustering algorithm to sequences of each block (Sklearn, https://scikit-learn.org/) enforcing the notion of two clusters being present (*i.e.*, “fast” and “slow”), labeling the IKIs of each sequence based on their proximity to them (*Figure 4a*). The outcome of this step was a chunking pattern for each individual sequence (*Figure 4b*).

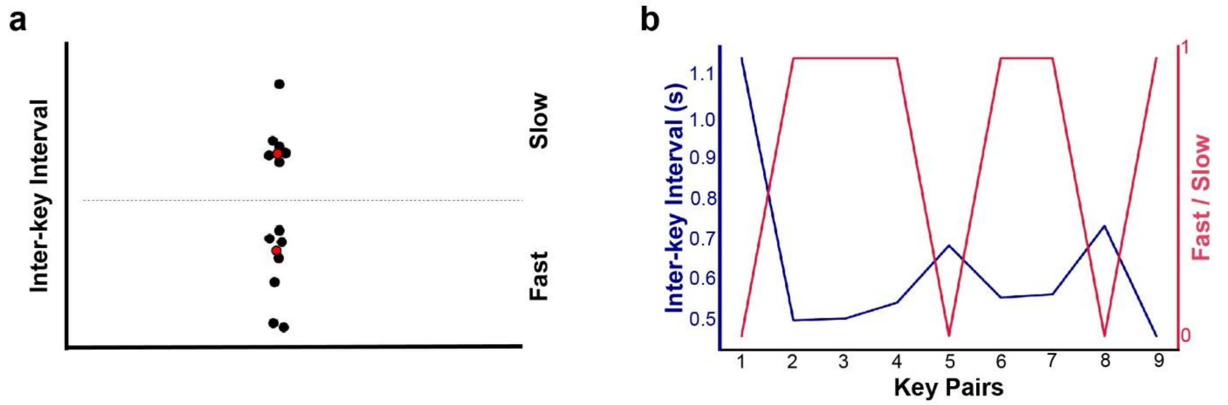


***Figure 4. Extraction of chunking patterns from IKIs.*** *a) We applied k-means clustering on IKIs from each block, defining two cluster centroids (i.e. “fast” and “slow”). b) After estimating the centroids, we labelled each IKI of the block as either “1” (i.e. fast) or “0” (i.e. slow), interpreting adjacent “1”’s as intervals belonging to the same motor chunk.*

Labelling each sequence in this way results in many different patterns. *Figure 5a* shows an example of such variability, in which each histogram bin corresponds to a different sequence pattern. As sequences are binary (i.e. consist of ones and zeros), we converted them to decimal to represent them as a single number.

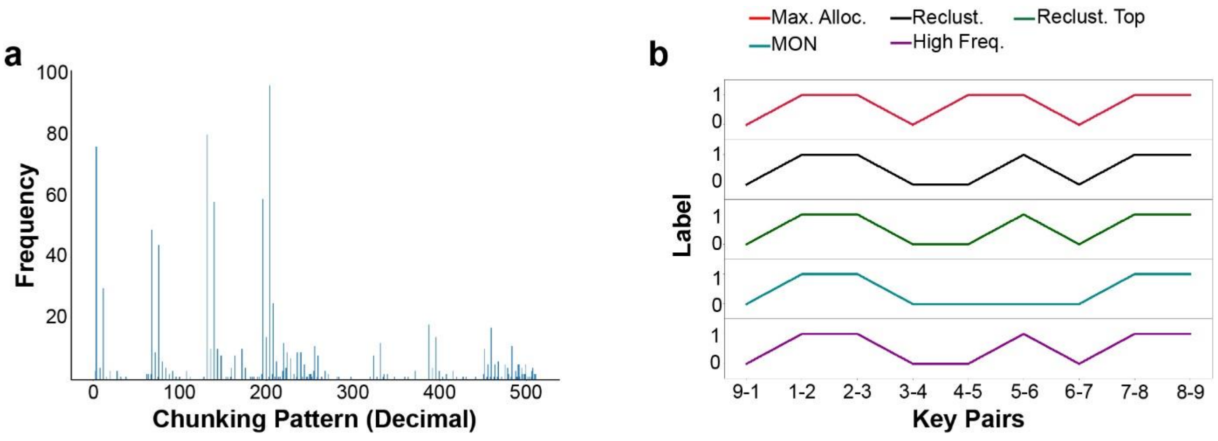


***Figure 5. Chunking patterns generated by a single subject.*** *a)Binary chunking patterns (e.g. [011011011]) are represented in decimal form (e.g. to* $(0)2^8+(1)2^7+(1)2^6+(0)2^5+(1)2^4+(1)2^3+(0)2^2+(1)2^1+(1)2^0=219$*). b) Output from each criterion for pattern selection.*

Because each participant generates multiple sequences, we defined the criteria described in the main text to determine a single pattern for each participant, on each day of training. *Figure5b* shows the output from the five criteria for a single participant on the first day of training. As some patterns are slightly different from one criterion to the next, we perform a majority vote using the five criteria. In this example, the resulting pattern would be [0 1 1 0 0 1 0 1 1].

### SICI measurements

In the main text, we mention that we used TMS to measure intra-cortical inhibition at rest in all participants of the second experiment. We applied the SICI paradigm before and after the first training day, to quantify the interneuronal GABAergic inhibition within the primary motor cortex, directly involved in the learning and execution of the motor sequence. We repeated these measurements on the fifth and sixtieth days. We show the group results below.

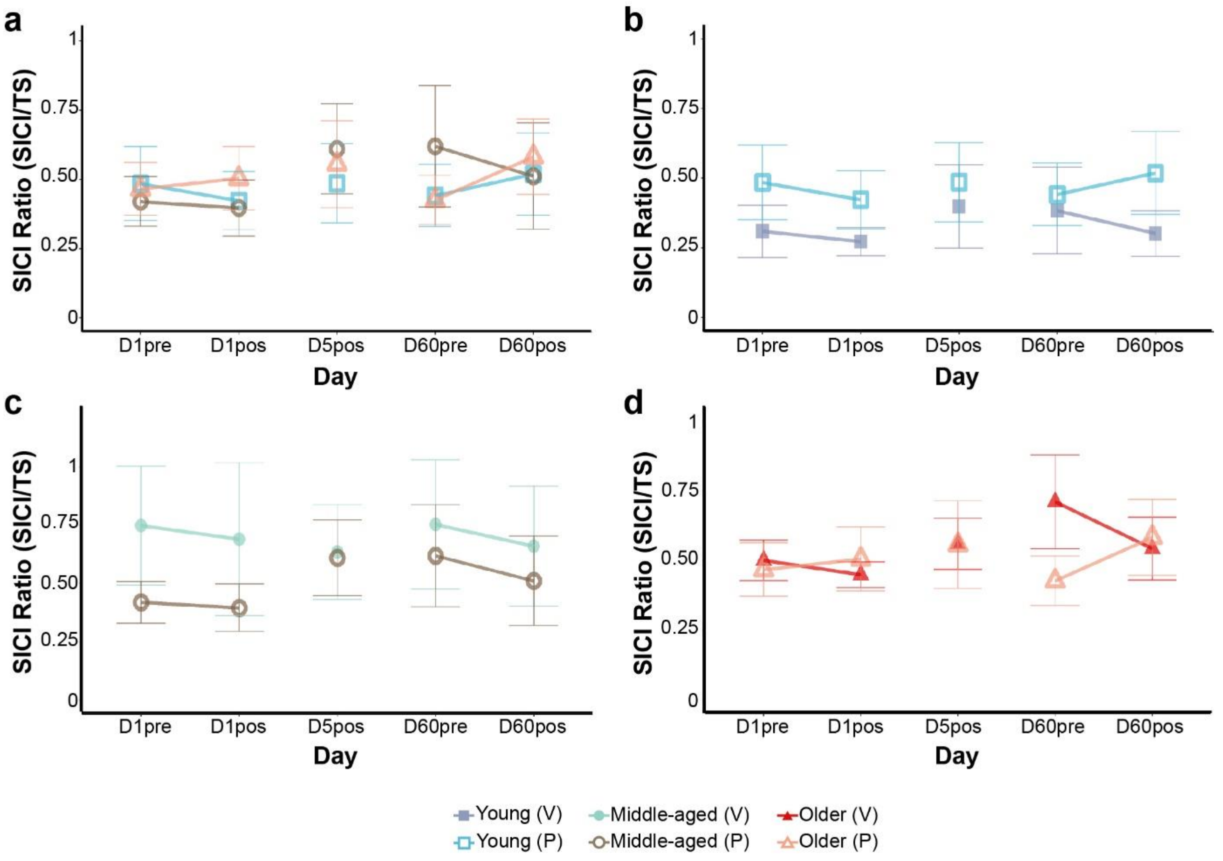


***Figure 6. SICI measurements at rest in participants of the second experiment.*** *Inhibitory efficiency, quantified as the ratio of the amplitude resulting from the SICI paradigm divided by the amplitude of the test stimulus (TS). Each marker corresponds to the average ratio per group, with its corresponding standard error.* ***a)*** *SICI at rest in all groups receiving placebo stimulation. Panels b to d show SICI measurements at rest in young (**b**), middle-aged (**c**) and older (**d**) adults receiving verum (V) and placebo (P) stimulation.*

**Table 1. Statistical tests run on behavioral data of the first experiment.**

| | | | | | ANOVA | | | | | PostHoc tests | | | | | | | |
|---|---|---|---|---|---|---|---|---|---|---|---|---|---|---|---|---|---|
| **Aspect** | **Model** | **Dependent** | **Independent** | **Random** | **ANOVA param.** | **DF num.** | **DF den.** | **F** | **p** | **Level** | **Contrast** | **Estimate** | **DF** | **p** | **d** | **CI** | |
| Baseline | LM | Score | AGE | NA | AGE | 2 | | 9.796767 | 0.000264 | B0 | Young - Middle | 1.642424 | 49 | 0.310889 | 0.494407 | -0.19425 | 1.183064 |
| | | | | | | | | | | | Young - Older | 4.909091 | 49 | 0.000163 | 1.477748 | 0.674939 | 2.280557 |
| | | | | | | | | | | | Middle - Older | 3.266667 | 49 | 0.025719 | 0.983341 | 0.19382 | 1.772861 |
| Baseline Accuracy | LM | Per. Correct | AGE | NA | AGE | 2 | | 0.946997 | 0.394888 | | | | | | | | |
| Block 1 Accuracy | LM | Per. Correct | AGE | NA | AGE | 2 | | 1.545493 | 0.223419 | | | | | | | | |
| Training | LMER | Score | AGE,DAY | (1 + 1\|ID) | AGE | 2 | 49 | 22.96936 | 9.17E-08 | day 1 | Young - Middle | 4.49989 | 55.33501 | 0.085361 | 1.092034 | 0.033397 | 2.15067 |
| | | | | | DAY | 4 | 1496 | 309.6215 | 3.95E-194 | | Young - Older | 6.203192 | 55.33501 | 0.011355 | 1.505391 | 0.403381 | 2.607401 |
| | | | | | AGE:DAY | 8 | 1496 | 30.80038 | 6.01E-45 | | Middle - Older | 1.703302 | 55.33501 | 0.732914 | 0.413357 | -0.69293 | 1.519643 |
| | | | | | | | | | | day 2 | Young - Middle | 8.07949 | 55.33501 | 0.000766 | 1.960731 | 0.797897 | 3.123566 |
| | | | | | | | | | | | Young - Older | 11.78042 | 55.33501 | 1.53E-06 | 2.858874 | 1.543574 | 4.174174 |
| | | | | | | | | | | | Middle - Older | 3.700933 | 55.33501 | 0.238972 | 0.898143 | -0.23295 | 2.029232 |
| | | | | | | | | | | day 3 | Young - Middle | 10.21638 | 55.33501 | 2.34E-05 | 2.479312 | 1.233054 | 3.72557 |
| | | | | | | | | | | | Young - Older | 14.85153 | 55.33501 | 5.92E-09 | 3.604172 | 2.137729 | 5.070615 |
| | | | | | | | | | | | Middle - Older | 4.635155 | 55.33501 | 0.109952 | 1.12486 | -0.02379 | 2.27351 |
| | | | | | | | | | | day 4 | Young - Middle | 10.08712 | 55.33501 | 2.92E-05 | 2.447944 | 1.20711 | 3.688778 |
| | | | | | | | | | | | Young - Older | 15.63477 | 55.33501 | 1.42E-09 | 3.794247 | 2.286533 | 5.30196 |
| | | | | | | | | | | | Middle - Older | 5.547643 | 55.33501 | 0.044959 | 1.346303 | 0.177046 | 2.515559 |
| | | | | | | | | | | day 5 | Young - Middle | 10.77967 | 55.33501 | 8.88E-06 | 2.616011 | 1.345595 | 3.886426 |
| | | | | | | | | | | | Young - Older | 17.07833 | 55.33501 | 1.08E-10 | 4.14457 | 2.558394 | 5.730745 |
| | | | | | | | | | | | Middle - Older | 6.298658 | 55.33501 | 0.019675 | 1.528559 | 0.339907 | 2.717211 |
| Online Learning | LMER | ΔScore | AGE,DAY | (1 + 1\|ID) | AGE | 2 | 48.98015 | 0.816388 | 0.447949 | day 1 | Young - Middle | 5.726304 | 244.9911 | 0.005344 | 1.05351 | 0.325554 | 1.781465 |
| | | | | | DAY | 4 | 195.9789 | 8.045557 | 5.02E-06 | | Young - Older | 9.151834 | 244.9911 | 2.96E-06 | 1.683729 | 0.861894 | 2.505564 |
| | | | | | AGE:DAY | 8 | 195.9789 | 4.486743 | 5.10E-05 | | Middle - Older | 3.42553 | 244.9911 | 0.198532 | 0.630219 | -0.11299 | 1.373425 |
| | | | | | | | | | | day 2 | Young - Middle | -3.49969 | 244.9911 | 0.135206 | -0.64386 | -1.33036 | 0.042632 |
| | | | | | | | | | | | Young - Older | -1.24583 | 244.9911 | 0.773348 | -0.22921 | -0.89309 | 0.434679 |
| | | | | | | | | | | | Middle - Older | 2.253854 | 244.9911 | 0.494159 | 0.414658 | -0.31566 | 1.144971 |
| | | | | | | | | | | day 3 | Young - Middle | -0.10921 | 244.9911 | 0.998023 | -0.02009 | -0.68066 | 0.640471 |
| | | | | | | | | | | | Young - Older | 0.917291 | 244.9911 | 0.869821 | 0.168761 | -0.49359 | 0.831115 |
| | | | | | | | | | | | Middle - Older | 1.026504 | 244.9911 | 0.863415 | 0.188853 | -0.53355 | 0.911255 |
| | | | | | | | | | | day 4 | Young - Middle | 0.925097 | 244.9911 | 0.867752 | 0.170197 | -0.49219 | 0.832582 |
| | | | | | | | | | | | Young - Older | -0.51353 | 244.9911 | 0.957198 | -0.09448 | -0.75559 | 0.566629 |
| | | | | | | | | | | | Middle - Older | -1.43863 | 244.9911 | 0.74966 | -0.26468 | -0.98908 | 0.459731 |
| | | | | | | | | | | day 5 | Young - Middle | -3.75057 | 244.9911 | 0.10096 | -0.69002 | -1.35066 | -0.02938 |
| | | | | | | | | | | | Young - Older | -3.85119 | 244.9911 | 0.089324 | -0.70853 | -1.36928 | -0.04778 |
| | | | | | | | | | | | Middle - Older | -0.10062 | 244.9911 | 0.998588 | -0.01851 | -0.73885 | 0.701825 |
| Online Slope | LM | Slope | AGE,DAY | NA | AGE | 2 | | 0.467166 | 0.627333 | day 1 | Young - Middle | 0.88315 | 245 | 0.008888 | 0.997667 | 0.277289 | 1.718045 |
| | | | | | DAY | 4 | | 12.64101 | 2.29E-09 | | Young - Older | 1.599765 | 245 | 4.78E-07 | 1.807205 | 0.964321 | 2.650089 |
| | | | | | AGE:DAY | 8 | | 4.776151 | 1.80E-05 | | Middle - Older | 0.716615 | 245 | 0.07031 | 0.809538 | 0.052857 | 1.566218 |
| | | | | | | | | | | day 2 | Young - Middle | -0.4177 | 245 | 0.33772 | -0.47186 | -1.14549 | 0.201768 |
| | | | | | | | | | | | Young - Older | -0.29685 | 245 | 0.576616 | -0.33535 | -1.00204 | 0.331348 |
| | | | | | | | | | | | Middle - Older | 0.120844 | 245 | 0.925878 | 0.136514 | -0.58381 | 0.856836 |
| | | | | | | | | | | day 3 | Young - Middle | 0.227412 | 245 | 0.723468 | 0.256901 | -0.40685 | 0.92065 |
| | | | | | | | | | | | Young - Older | 0.123662 | 245 | 0.908566 | 0.139697 | -0.52109 | 0.800486 |
| | | | | | | | | | | | Middle - Older | -0.10375 | 245 | 0.944806 | -0.1172 | -0.83724 | 0.602831 |
| | | | | | | | | | | day 4 | Young - Middle | 0.123356 | 245 | 0.908996 | 0.139351 | -0.52143 | 0.800134 |
| | | | | | | | | | | | Young - Older | -0.09555 | 245 | 0.944342 | -0.10794 | -0.76823 | 0.552345 |
| | | | | | | | | | | | Middle - Older | -0.21891 | 245 | 0.776961 | -0.24729 | -0.9701 | 0.475514 |
| | | | | | | | | | | day 5 | Young - Middle | -0.45778 | 245 | 0.272071 | -0.51714 | -1.17794 | 0.14366 |
| | | | | | | | | | | | Young - Older | -0.7003 | 245 | 0.049438 | -0.7911 | -1.45178 | -0.13042 |
| | | | | | | | | | | | Middle - Older | -0.24251 | 245 | 0.733748 | -0.27396 | -0.99758 | 0.449656 |
| | | | | | | | | | | Young | day 1 - day 2 | 1.724773 | 245 | 5.52E-09 | 1.948423 | 1.128125 | 2.768721 |
| | | | | | | | | | | | day 1 - day 3 | 1.459651 | 245 | 1.11E-06 | 1.648923 | 0.886021 | 2.411824 |
| | | | | | | | | | | | day 1 - day 4 | 1.656072 | 245 | 2.31E-08 | 1.870814 | 1.065898 | 2.675729 |
| | | | | | | | | | | | day 1 - day 5 | 2.18587 | 245 | 1.93E-13 | 2.469309 | 1.675861 | 3.262757 |
| | | | | | | | | | | | day 2 - day 3 | -0.26512 | 245 | 0.85819 | -0.2995 | -0.89972 | 0.30072 |
| | | | | | | | | | | | day 2 - day 4 | -0.0687 | 245 | 0.999028 | -0.07761 | -0.67192 | 0.516703 |
| | | | | | | | | | | | day 2 - day 5 | 0.461096 | 245 | 0.418962 | 0.520886 | -0.07432 | 1.116095 |
| | | | | | | | | | | | day 3 - day 4 | 0.196421 | 245 | 0.947895 | 0.221891 | -0.37548 | 0.819262 |
| | | | | | | | | | | | day 3 - day 5 | 0.726218 | 245 | 0.05377 | 0.820386 | 0.22462 | 1.416152 |
| | | | | | | | | | | | day 4 - day 5 | 0.529797 | 245 | 0.276395 | 0.598496 | 0.004363 | 1.192628 |
| | | | | | | | | | | Middle | day 1 - day 2 | 0.423926 | 245 | 0.68433 | 0.478896 | -0.25366 | 1.21145 |
| | | | | | | | | | | | day 1 - day 3 | 0.803914 | 245 | 0.096898 | 0.908156 | 0.142093 | 1.67422 |
| | | | | | | | | | | | day 1 - day 4 | 0.896278 | 245 | 0.046791 | 1.012498 | 0.235481 | 1.789514 |
| | | | | | | | | | | | day 1 - day 5 | 0.844939 | 245 | 0.070924 | 0.954501 | 0.183701 | 1.725302 |
| | | | | | | | | | | | day 2 - day 3 | 0.379988 | 245 | 0.765347 | 0.429261 | -0.30069 | 1.159215 |
| | | | | | | | | | | | day 2 - day 4 | 0.472353 | 245 | 0.588596 | 0.533602 | -0.20213 | 1.269338 |
| | | | | | | | | | | | day 2 - day 5 | 0.421013 | 245 | 0.689924 | 0.475606 | -0.25677 | 1.207979 |
| | | | | | | | | | | | day 3 - day 4 | 0.092365 | 245 | 0.998535 | 0.104341 | -0.61553 | 0.82421 |
| | | | | | | | | | | | day 3 - day 5 | 0.041025 | 245 | 0.999941 | 0.046345 | -0.67301 | 0.765701 |
| | | | | | | | | | | | day 4 - day 5 | -0.05134 | 245 | 0.999856 | -0.058 | -0.77742 | 0.661431 |
| | | | | | | | | | | Older | day 1 - day 2 | -0.17185 | 245 | 0.984018 | -0.19413 | -0.91556 | 0.527309 |
| | | | | | | | | | | | day 1 - day 3 | -0.01645 | 245 | 0.999998 | -0.01859 | -0.73784 | 0.700665 |
| | | | | | | | | | | | day 1 - day 4 | -0.03924 | 245 | 0.999951 | -0.04433 | -0.76368 | 0.675012 |
| | | | | | | | | | | | day 1 - day 5 | -0.11419 | 245 | 0.996646 | -0.129 | -0.8492 | 0.591207 |
| | | | | | | | | | | | day 2 - day 3 | 0.155393 | 245 | 0.989055 | 0.175542 | -0.54549 | 0.896578 |
| | | | | | | | | | | | day 2 - day 4 | 0.1326 | 245 | 0.994024 | 0.149794 | -0.57075 | 0.870339 |
| | | | | | | | | | | | day 2 - day 5 | 0.057654 | 245 | 0.999772 | 0.06513 | -0.65435 | 0.784609 |
| | | | | | | | | | | | day 3 - day 4 | -0.02279 | 245 | 0.999994 | -0.02575 | -0.74502 | 0.693521 |
| | | | | | | | | | | | day 3 - day 5 | -0.09774 | 245 | 0.998171 | -0.11041 | -0.83036 | 0.609532 |
| | | | | | | | | | | | day 4 - day 5 | -0.07495 | 245 | 0.999356 | -0.08466 | -0.80432 | 0.634986 |
| Offline Learning | LM | ΔScore | AGE,NIGHT | NA | AGE | 2 | | 8.27757 | 0.000354 | AGE | Young - Middle | 2.536574 | 196 | 0.002843 | 0.559672 | 0.21987 | 0.899474 |
| | | | | | NIGHT | 3 | | 0.079903 | 0.970857 | AGE | Young - Older | 2.637295 | 196 | 0.001809 | 0.581895 | 0.243562 | 0.920229 |
| | | | | | AGE:NIGHT | 6 | | 1.753384 | 0.110611 | AGE | Middle - Older | 0.100721 | 196 | 0.991865 | 0.022223 | -0.3379 | 0.382343 |
| Speed | LMER | Seq. Number | AGE,DAY | (1 + 1\|ID) | AGE | 2 | 48.99997 | 28.30728 | 6.74E-09 | day 1 | Young - Middle | 2.769697 | 53.16693 | 0.246343 | 0.989968 | -0.27039 | 2.250327 |
| | | | | | DAY | 4 | 1496 | 734.2932 | 0 | | Young - Older | 4.869697 | 53.16693 | 0.016918 | 1.740568 | 0.41101 | 3.070125 |
| | | | | | AGE:DAY | 8 | 1496 | 84.18678 | 3.80E-115 | | Middle - Older | 2.1 | 53.16693 | 0.502437 | 0.750599 | -0.60453 | 2.105733 |
| | | | | | | | | | | day 2 | Young - Middle | 6.169192 | 53.16693 | 0.001945 | 2.205044 | 0.816527 | 3.59356 |
| | | | | | | | | | | | Young - Older | 10.72475 | 53.16693 | 1.97E-07 | 3.833328 | 2.163674 | 5.502982 |
| | | | | | | | | | | | Middle - Older | 4.555556 | 53.16693 | 0.046412 | 1.628284 | 0.207383 | 3.049186 |
| | | | | | | | | | | day 3 | Young - Middle | 8.530303 | 53.16693 | 2.03E-05 | 3.048972 | 1.527227 | 4.570716 |
| | | | | | | | | | | | Young - Older | 13.49697 | 53.16693 | 4.85E-10 | 4.824199 | 2.943273 | 6.705125 |
| | | | | | | | | | | | Middle - Older | 4.966667 | 53.16693 | 0.027159 | 1.775227 | 0.339019 | 3.211436 |
| | | | | | | | | | | day 4 | Young - Middle | 9.204545 | 53.16693 | 5.01E-06 | 3.289965 | 1.724945 | 4.854986 |
| | | | | | | | | | | | Young - Older | 15.21566 | 53.16693 | 1.07E-11 | 5.438506 | 3.416337 | 7.460676 |
| | | | | | | | | | | | Middle - Older | 6.011111 | 53.16693 | 0.00604 | 2.148541 | 0.668412 | 3.62867 |
| | | | | | | | | | | day 5 | Young - Middle | 10.33232 | 53.16693 | 4.57E-07 | 3.693065 | 2.051288 | 5.334842 |
| | | | | | | | | | | | Young - Older | 17.34343 | 53.16693 | 0 | 6.199034 | 3.993788 | 8.40428 |
| | | | | | | | | | | | Middle - Older | 7.011111 | 53.16693 | 0.001218 | 2.50597 | 0.977487 | 4.034452 |
| | | | | | | | | | | Week | Young - Middle | 7.401212 | 49 | 0.000161 | 2.645403 | 1.209166 | 4.081639 |
| | | | | | | | | | | | Young - Older | 12.3301 | 49 | 5.52E-09 | 4.407127 | 2.631304 | 6.18295 |
| | | | | | | | | | | | Middle - Older | 4.928889 | 49 | 0.025425 | 1.761724 | 0.349317 | 3.174132 |
| | | | | | | | | | | Young | day 1 - day 2 | -7.31061 | 1496 | 7.44E-13 | -2.61302 | -3.42419 | -1.80184 |
| | | | | | | | | | | | day 1 - day 3 | -11.8939 | 1496 | 7.44E-13 | -4.25123 | -5.53237 | -2.97009 |
| | | | | | | | | | | | day 1 - day 4 | -14.7348 | 1496 | 7.44E-13 | -5.26665 | -6.8435 | -3.68981 |
| | | | | | | | | | | | day 1 - day 5 | -17.8182 | 1496 | 7.44E-13 | -6.36872 | -8.26813 | -4.46932 |
| | | | | | | | | | | | day 2 - day 3 | -4.58333 | 1496 | 7.44E-13 | -1.63821 | -2.18193 | -1.0945 |
| | | | | | | | | | | | day 2 - day 4 | -7.42424 | 1496 | 7.44E-13 | -2.65363 | -3.47626 | -1.83101 |
| | | | | | | | | | | | day 2 - day 5 | -10.5076 | 1496 | 7.44E-13 | -3.7557 | -4.89341 | -2.618 |
| | | | | | | | | | | | day 3 - day 4 | -2.84091 | 1496 | 8.04E-13 | -1.01542 | -1.40415 | -0.6267 |
| | | | | | | | | | | | day 3 - day 5 | -5.92424 | 1496 | 7.44E-13 | -2.11749 | -2.79056 | -1.44442 |
| | | | | | | | | | | | day 4 - day 5 | -3.08333 | 1496 | 7.86E-13 | -1.10207 | -1.51091 | -0.69323 |
| | | | | | | | | | | | day 1 - day 2 | -3.91111 | 1496 | 7.93E-13 | -1.39794 | -1.90811 | -0.88777 |

| | | | | | | | | | | | | | | | | | |
|---|---|---|---|---|---|---|---|---|---|---|---|---|---|---|---|---|---|
| | | | | | | | | | | | day 1 - day 3 | -6.13333 | 1496 | 7.44E-13 | -2.19223 | -2.90611 | -1.47834 |
| | | | | | | | | | | | day 1 - day 4 | -8.3 | 1496 | 7.44E-13 | -2.96665 | -3.89347 | -2.03984 |
| | | | | | | | | | | | day 1 - day 5 | -10.2556 | 1496 | 7.44E-13 | -3.66563 | -4.79006 | -2.54119 |
| | | | | | | | | | | Middle | day 2 - day 3 | -2.22222 | 1496 | 1.14E-06 | -0.79429 | -1.17449 | -0.41408 |
| | | | | | | | | | | | day 2 - day 4 | -4.38889 | 1496 | 7.93E-13 | -1.56871 | -2.1206 | -1.01683 |
| | | | | | | | | | | | day 2 - day 5 | -6.34444 | 1496 | 7.44E-13 | -2.26768 | -3.00189 | -1.53348 |
| | | | | | | | | | | | day 3 - day 4 | -2.16667 | 1496 | 2.31E-06 | -0.77443 | -1.15103 | -0.39782 |
| | | | | | | | | | | | day 3 - day 5 | -4.12222 | 1496 | 7.90E-13 | -1.4734 | -2.00181 | -0.94499 |
| | | | | | | | | | | | day 4 - day 5 | -1.95556 | 1496 | 2.95E-05 | -0.69897 | -1.06244 | -0.3355 |
| | | | | | | | | | | | day 1 - day 2 | -1.45556 | 1496 | 0.00452 | -0.52026 | -0.85649 | -0.18402 |
| | | | | | | | | | | | day 1 - day 3 | -3.26667 | 1496 | 8.84E-13 | -1.1676 | -1.62433 | -0.71087 |
| | | | | | | | | | | | day 1 - day 4 | -4.38889 | 1496 | 7.93E-13 | -1.56871 | -2.1206 | -1.01683 |
| | | | | | | | | | | | day 1 - day 5 | -5.34444 | 1496 | 7.44E-13 | -1.91026 | -2.54938 | -1.27113 |
| | | | | | | | | | | Older | day 2 - day 3 | -1.81111 | 1496 | 0.000146 | -0.64734 | -1.00235 | -0.29233 |
| | | | | | | | | | | | day 2 - day 4 | -2.93333 | 1496 | 3.14E-11 | -1.04846 | -1.47917 | -0.61775 |
| | | | | | | | | | | | day 2 - day 5 | -3.88889 | 1496 | 7.93E-13 | -1.39 | -1.89827 | -0.88173 |
| | | | | | | | | | | | day 3 - day 4 | -1.12222 | 1496 | 0.055781 | -0.40111 | -0.72276 | -0.07947 |
| | | | | | | | | | | | day 3 - day 5 | -2.07778 | 1496 | 6.96E-06 | -0.74266 | -1.11362 | -0.37169 |
| | | | | | | | | | | | day 4 - day 5 | -0.95556 | 1496 | 0.148234 | -0.34154 | -0.65712 | -0.02597 |
| | | | | | AGE | 2 | 72.52015 | 5.837485 | 0.00446 | | Young - Middle | -0.00122 | 75.56009 | 0.998793 | -0.01294 | -0.56385 | 0.537963 |
| | | | | | DAY | 4 | 1481 | 10.23586 | 3.55E-08 | day 1 | Young - Older | 0.06209 | 75.56009 | 0.050087 | 0.661015 | 0.076918 | 1.245112 |
| | | | | | BLOCK | 1 | 1481 | 19.87458 | 8.89E-06 | | Middle - Older | 0.063306 | 75.56009 | 0.071861 | 0.673958 | 0.041443 | 1.306473 |
| | | | | | AGE:DAY | 8 | 1481 | 2.510816 | 0.010367 | | Young - Middle | 0.019482 | 75.69673 | 0.734812 | 0.207406 | -0.34708 | 0.76189 |
| | | | | | AGE:BLOCK | 2 | 1481 | 3.748352 | 0.02378 | day 2 | Young - Older | 0.077628 | 75.69673 | 0.010492 | 0.82643 | 0.224228 | 1.428631 |
| | | | | | DAY:BLOCK | 4 | 1481 | 1.115982 | 0.34729 | | Middle - Older | 0.058146 | 75.69673 | 0.107076 | 0.619024 | -0.00887 | 1.246922 |
| | | | | | AGE:DAY:BLOCK | 8 | 1481 | 3.131654 | 0.00162 | | Young - Middle | 0.014007 | 75.8786 | 0.852564 | 0.149124 | -0.40405 | 0.702301 |
| | | | | | | | | | | day 3 | Young - Older | 0.076204 | 75.8786 | 0.012299 | 0.81127 | 0.210566 | 1.411974 |
| | | | | | | | | | | | Middle - Older | 0.062197 | 75.8786 | 0.078833 | 0.662146 | 0.030144 | 1.294149 |
| | | | | | | | | | | | Young - Middle | -0.007 | 75.69673 | 0.960785 | -0.07456 | -0.62612 | 0.477003 |
| | | | | | | | | | | day 4 | Young - Older | 0.040223 | 75.69673 | 0.274792 | 0.428218 | -0.13708 | 0.993511 |
| | | | | | | | | | | | Middle - Older | 0.047227 | 75.69673 | 0.22484 | 0.502778 | -0.1161 | 1.121654 |
| | | | | | | | | | | | Young - Middle | -0.01204 | 75.56009 | 0.888484 | -0.12821 | -0.68039 | 0.423968 |
| | | | | | | | | | | day 5 | Young - Older | 0.021121 | 75.56009 | 0.69616 | 0.224859 | -0.32998 | 0.779696 |
| | | | | | | | | | | | Middle - Older | 0.033164 | 75.56009 | 0.474273 | 0.353069 | -0.25656 | 0.962701 |
| | | | | | | | | | | | day 1 - day 2 | -0.0376 | 1481 | 0.010396 | -0.40034 | -0.67249 | -0.1282 |
| | | | | | | | | | | | day 1 - day 3 | -0.05016 | 1481 | 0.000156 | -0.53405 | -0.82557 | -0.24252 |
| | | | | | | | | | | | day 1 - day 4 | -0.05014 | 1481 | 0.000153 | -0.53383 | -0.82507 | -0.24259 |
| | | | | | | | | | | | day 1 - day 5 | -0.04937 | 1481 | 0.000202 | -0.52555 | -0.81529 | -0.2358 |
| | | | | | | | | | | Young | day 2 - day 3 | -0.01256 | 1481 | 0.815519 | -0.1337 | -0.38278 | 0.115376 |
| | | | | | | | | | | | day 2 - day 4 | -0.01254 | 1481 | 0.815724 | -0.13349 | -0.38226 | 0.115281 |
| | | | | | | | | | | | day 2 - day 5 | -0.01176 | 1481 | 0.848043 | -0.1252 | -0.37338 | 0.122974 |
| Accuracy | LMER | Per. Correct | AGE,DAY,BLOCK | (1 + 1\|ID) | | | | | | | day 3 - day 4 | 2.02E-05 | 1481 | 1 | 0.000215 | -0.24575 | 0.246179 |
| | | | | | | | | | | | day 3 - day 5 | 0.000798 | 1481 | 0.999995 | 0.008499 | -0.23726 | 0.254255 |
| | | | | | | | | | | | day 4 - day 5 | 0.000778 | 1481 | 0.999995 | 0.008284 | -0.23717 | 0.253736 |
| | | | | | | | | | | | day 1 - day 2 | -0.01691 | 1481 | 0.747838 | -0.17999 | -0.4819 | 0.121911 |
| | | | | | | | | | | | day 1 - day 3 | -0.03494 | 1481 | 0.09349 | -0.37198 | -0.68901 | -0.05495 |
| | | | | | | | | | | | day 1 - day 4 | -0.05593 | 1481 | 0.000661 | -0.59545 | -0.94031 | -0.25058 |
| | | | | | | | | | | | day 1 - day 5 | -0.06019 | 1481 | 0.000178 | -0.64081 | -0.99239 | -0.28924 |
| | | | | | | | | | | Middle | day 2 - day 3 | -0.01803 | 1481 | 0.701327 | -0.19198 | -0.49515 | 0.111182 |
| | | | | | | | | | | | day 2 - day 4 | -0.03902 | 1481 | 0.04351 | -0.41545 | -0.73701 | -0.0939 |
| | | | | | | | | | | | day 2 - day 5 | -0.04329 | 1481 | 0.017498 | -0.46082 | -0.78742 | -0.13422 |
| | | | | | | | | | | | day 3 - day 4 | -0.02099 | 1481 | 0.566373 | -0.22347 | -0.52849 | 0.081553 |
| | | | | | | | | | | | day 3 - day 5 | -0.02525 | 1481 | 0.37431 | -0.26883 | -0.57674 | 0.03907 |
| | | | | | | | | | | | day 4 - day 5 | -0.00426 | 1481 | 0.998144 | -0.04537 | -0.34291 | 0.252174 |
| | | | | | | | | | | | day 1 - day 2 | -0.02207 | 1481 | 0.514196 | -0.23493 | -0.54007 | 0.070216 |
| | | | | | | | | | | | day 1 - day 3 | -0.03605 | 1481 | 0.076765 | -0.38379 | -0.70203 | -0.06555 |
| | | | | | | | | | | | day 1 - day 4 | -0.07201 | 1481 | 3.14E-06 | -0.76663 | -1.13951 | -0.39374 |
| | | | | | | | | | | | day 1 - day 5 | -0.09033 | 1481 | 1.50E-09 | -0.9617 | -1.37153 | -0.55188 |
| | | | | | | | | | | Older | day 2 - day 3 | -0.01398 | 1481 | 0.857507 | -0.14886 | -0.44993 | 0.152206 |
| | | | | | | | | | | | day 2 - day 4 | -0.04994 | 1481 | 0.003513 | -0.5317 | -0.8677 | -0.1957 |
| | | | | | | | | | | | day 2 - day 5 | -0.06827 | 1481 | 1.22E-05 | -0.72677 | -1.09271 | -0.36084 |
| | | | | | | | | | | | day 3 - day 4 | -0.03596 | 1481 | 0.078489 | -0.38284 | -0.70123 | -0.06445 |
| | | | | | | | | | | | day 3 - day 5 | -0.05428 | 1481 | 0.001083 | -0.57791 | -0.92052 | -0.23531 |
| | | | | | | | | | | | day 4 - day 5 | -0.01832 | 1481 | 0.686775 | -0.19507 | -0.49779 | 0.107638 |
| | | | | | AGE | 2 | 48.99998 | 27.97434 | 7.87E-09 | | Young - Middle | 10.0237 | 57.78333 | 0.000423 | 2.222246 | 0.946917 | 3.497575 |
| | | | | | DAY | 2 | 410 | 10.36045 | 4.08E-05 | day 5 | Young - Older | 15.71304 | 57.78333 | 9.82E-08 | 3.483569 | 1.982418 | 4.98472 |
| | | | | | AGE:DAY | 4 | 410 | 1.726828 | 0.143117 | | Middle - Older | 5.689342 | 57.78333 | 0.095219 | 1.261322 | 0.011863 | 2.510782 |
| | | | | | | | | | | | Young - Middle | 10.0269 | 57.78333 | 0.000421 | 2.222955 | 0.947518 | 3.498391 |
| | | | | | | | | | | day 10 | Young - Older | 17.54548 | 57.78333 | 5.63E-09 | 3.88982 | 2.304128 | 5.475511 |
| | | | | | | | | | | | Middle - Older | 7.518588 | 57.78333 | 0.018852 | 1.666865 | 0.376669 | 2.95706 |
| | | | | | | | | | | | Young - Middle | 11.75296 | 57.78333 | 3.82E-05 | 2.605622 | 1.268568 | 3.942676 |
| | | | | | | | | | | day 60 | Young - Older | 18.45429 | 57.78333 | 1.37E-09 | 4.0913 | 2.462034 | 5.720567 |
| | | | | | | | | | | | Middle - Older | 6.701326 | 57.78333 | 0.040554 | 1.485679 | 0.214914 | 2.756443 |
| | | | | | | | | | | | Young - Middle | 10.60119 | 49 | 0.000129 | 2.350274 | 1.087538 | 3.613011 |
| FU Scores | LMER | Score | AGE,DAY | (1 + 1\|ID) | | | | | | All | Young - Older | 17.2376 | 49 | 7.03E-09 | 3.821563 | 2.275052 | 5.368074 |
| | | | | | | | | | | | Middle - Older | 6.636418 | 49 | 0.034474 | 1.471289 | 0.242662 | 2.699915 |
| | | | | | | | | | | | day 5 - day 10 | -1.92449 | 410 | 0.033209 | -0.43706 | -0.80866 | -0.06547 |
| | | | | | | | | | | Young | day 5 - day 60 | -0.50385 | 410 | 0.788305 | -0.11443 | -0.46454 | 0.235687 |
| | | | | | | | | | | | day 10 - day 60 | 1.42064 | 410 | 0.153786 | 0.322635 | -0.03862 | 0.683894 |
| | | | | | | | | | | | day 5 - day 10 | -1.9213 | 410 | 0.09745 | -0.43634 | -0.87758 | 0.004909 |
| | | | | | | | | | | Middle | day 5 - day 60 | 1.225407 | 410 | 0.38479 | 0.278297 | -0.15166 | 0.708249 |
| | | | | | | | | | | | day 10 - day 60 | 3.146705 | 410 | 0.00221 | 0.714634 | 0.242825 | 1.186443 |
| | | | | | | | | | | | day 5 - day 10 | -0.09205 | 410 | 0.994594 | -0.02091 | -0.44298 | 0.401173 |
| | | | | | | | | | | Older | day 5 - day 60 | 2.237392 | 410 | 0.043171 | 0.508124 | 0.060234 | 0.956014 |
| | | | | | | | | | | | day 10 - day 60 | 2.329443 | 410 | 0.033326 | 0.529029 | 0.079036 | 0.979023 |
| | | | | | AGE | 2 | 48.99995 | 16.57653 | 3.17E-06 | | | | | | | | |
| Rand. Block Scores | LMER | Score | AGE, DAY | (1 + 1\|ID) | DAY | 5 | 245 | 1.233546 | 0.293915 | | | | | | | | |
| | | | | | AGE:DAY | 10 | 245 | 0.807684 | 0.621469 | | | | | | | | |

**Table 2. Statistical tests run on behavioral data from the placebo groups of the second experiment.**

| | | | | | ANOVA | | | | | PostHoc tests | | | | | | | |
|---|---|---|---|---|---|---|---|---|---|---|---|---|---|---|---|---|---|
| Aspect | Model | Dependent | Independent | Random | ANOVA param. | DF num. | DF den. | F | p | Level | Contrast | Estimate | DF | p | d | CI | |
| Baseline | LM | Score | AGE | NA | AGE | 2 | | 2.2342 | 0.128025 | | | | | | | | |
| Baseline Accuracy | LM | Per. Correct | AGE | NA | AGE | 2 | | 0.959892 | 0.396603 | | | | | | | | |
| Block 1 Accuracy | LM | Per. Correct | AGE | NA | AGE | 2 | | 7.07238 | 0.00368 | B1 | Young - Middle | -0.09889 | 25 | 0.315589 | -0.68169 | -1.65036 | 0.28698 |
| | | | | | | | | | | | Young - Older | 0.150828 | 25 | 0.08977 | 1.039668 | 0.018751 | 2.060585 |
| | | | | | | | | | | | Middle - Older | 0.249722 | 25 | 0.002633 | 1.721358 | 0.640295 | 2.802421 |
| Training | LMER | Score | AGE,DAY | (1 + 1\|ID) | AGE | 2 | 24.99991 | 12.6383 | 0.000161 | day 1 | Young - Middle | 2.079861 | 32.63027 | 0.467897 | 0.564413 | -0.4163 | 1.545122 |
| | | | | | DAY | 4 | 800 | 185.2798 | 2.44E-112 | | Young - Older | 2.503491 | 32.63027 | 0.355068 | 0.679373 | -0.33245 | 1.691193 |
| | | | | | AGE:DAY | 8 | 800 | 15.42003 | 3.32E-21 | | Middle - Older | 0.42363 | 32.63027 | 0.968203 | 0.114961 | -0.85163 | 1.081548 |
| | | | | | | | | | | day 2 | Young - Middle | 5.084451 | 32.63027 | 0.017414 | 1.37977 | 0.328791 | 2.430748 |
| | | | | | | | | | | | Young - Older | 6.968115 | 32.63027 | 0.001343 | 1.89094 | 0.748907 | 3.032973 |
| | | | | | | | | | | | Middle - Older | 1.883664 | 32.63027 | 0.534798 | 0.511171 | -0.46691 | 1.489247 |
| | | | | | | | | | | day 3 | Young - Middle | 7.55981 | 32.63027 | 0.00039 | 2.051509 | 0.906022 | 3.196995 |
| | | | | | | | | | | | Young - Older | 8.244156 | 32.63027 | 0.000179 | 2.23722 | 1.040148 | 3.434292 |
| | | | | | | | | | | | Middle - Older | 0.684346 | 32.63027 | 0.91927 | 0.185711 | -0.78187 | 1.153289 |
| | | | | | | | | | | day 4 | Young - Middle | 7.001167 | 32.63027 | 0.000962 | 1.89991 | 0.778219 | 3.021601 |
| | | | | | | | | | | | Young - Older | 9.812826 | 32.63027 | 1.40E-05 | 2.662911 | 1.389787 | 3.936034 |
| | | | | | | | | | | | Middle - Older | 2.811659 | 32.63027 | 0.256801 | 0.763001 | -0.22974 | 1.75574 |
| | | | | | | | | | | day 5 | Young - Middle | 9.204791 | 32.63027 | 2.56E-05 | 2.497908 | 1.275189 | 3.720627 |
| | | | | | | | | | | | Young - Older | 12.6187 | 32.63027 | 1.51E-07 | 3.424343 | 1.996659 | 4.852026 |
| | | | | | | | | | | | Middle - Older | 3.413913 | 32.63027 | 0.140464 | 0.926435 | -0.07875 | 1.931621 |
| Online learning D1 | LM | ΔScore | AGE | NA | AGE | 2 | | 3.21531 | 0.057192 | | | | | | | | |
| Online learning D2-D5 | LMER | ΔScore | AGE,DAY | (1 + 1\|ID) | AGE | 2 | 25 | 0.518973 | 0.601404 | | | | | | | | |
| | | | | | DAY | 3 | 75 | 2.0123 | 0.119427 | | | | | | | | |
| | | | | | AGE:DAY | 6 | 75 | 0.415053 | 0.866847 | | | | | | | | |
| Online slope D1 | LM | Slope | AGE | NA | AGE | 2 | | 3.057761 | 0.064869 | | | | | | | | |
| Online slope D2-D5 | LMER | Slope | AGE,DAY | (1 + 1\|ID) | AGE | 2 | 24.99995 | 0.825421 | 0.449639 | | | | | | | | |
| | | | | | DAY | 3 | 75.00003 | 4.078163 | 0.009714 | | | | | | | | |
| | | | | | AGE:DAY | 6 | 75.00004 | 1.332544 | 0.253411 | | | | | | | | |
| Offline Learning | LM | ΔScore | AGE,NIGHT | NA | AGE | 2 | | 4.129986 | 0.018906 | AGE | Young - Middle | 2.19457 | 100 | 0.117382 | 0.459559 | -0.00344 | 0.922554 |
| | | | | | NIGHT | 3 | | 0.620068 | 0.603626 | | Young - Older | 3.148837 | 100 | 0.016858 | 0.65939 | 0.191201 | 1.127578 |
| | | | | | AGE:NIGHT | 6 | | 0.378325 | 0.891285 | | Middle - Older | 0.954267 | 100 | 0.660507 | 0.199831 | -0.25969 | 0.65935 |
| Speed | LMER | Seq. Number | AGE,DAY | (1 + 1\|ID) | AGE | 2 | 25 | 13.7418 | 9.42E-05 | day 1 | Young - Middle | 2.133333 | 27.4524 | 0.388703 | 1.046805 | -0.5923 | 2.685907 |
| | | | | | DAY | 4 | 800 | 600.4129 | 3.67E-239 | | Young - Older | 2.962963 | 27.4524 | 0.186234 | 1.453895 | -0.25368 | 3.161472 |
| | | | | | AGE:DAY | 8 | 800 | 40.8779 | 7.82E-55 | | Middle - Older | 0.82963 | 27.4524 | 0.862775 | 0.407091 | -1.20588 | 2.020059 |
| | | | | | | | | | | day 2 | Young - Middle | 4.896296 | 27.4524 | 0.013082 | 2.402562 | 0.637909 | 4.167215 |
| | | | | | | | | | | | Young - Older | 6.611111 | 27.4524 | 0.001124 | 3.244004 | 1.324549 | 5.163459 |
| | | | | | | | | | | | Middle - Older | 1.714815 | 27.4524 | 0.538656 | 0.841442 | -0.78682 | 2.469699 |
| | | | | | | | | | | day 3 | Young - Middle | 6.72963 | 27.4524 | 0.000701 | 3.30216 | 1.409288 | 5.195032 |
| | | | | | | | | | | | Young - Older | 9.111111 | 27.4524 | 1.90E-05 | 4.470728 | 2.337861 | 6.603596 |
| | | | | | | | | | | | Middle - Older | 2.381481 | 27.4524 | 0.311299 | 1.168568 | -0.47804 | 2.815173 |
| | | | | | | | | | | day 4 | Young - Middle | 6.97963 | 27.4524 | 0.000462 | 3.424832 | 1.512145 | 5.33752 |
| | | | | | | | | | | | Young - Older | 10.25926 | 27.4524 | 2.95E-06 | 5.034113 | 2.789462 | 7.278763 |
| | | | | | | | | | | | Middle - Older | 3.27963 | 27.4524 | 0.119009 | 1.60928 | -0.07095 | 3.289512 |
| | | | | | | | | | | day 5 | Young - Middle | 8.214815 | 27.4524 | 5.81E-05 | 4.030925 | 2.013176 | 6.048674 |
| | | | | | | | | | | | Young - Older | 11.77778 | 27.4524 | 2.70E-07 | 5.779234 | 3.376166 | 8.182302 |
| | | | | | | | | | | | Middle - Older | 3.562963 | 27.4524 | 0.084 | 1.748309 | 0.055431 | 3.441187 |
| | | | | | | | | | | Young | day 1 - day 2 | -5.7963 | 800 | 0 | -2.84418 | -3.79017 | -1.89819 |
| | | | | | | | | | | | day 1 - day 3 | -9.62963 | 800 | 0 | -4.72516 | -6.20704 | -3.24328 |
| | | | | | | | | | | | day 1 - day 4 | -12.1296 | 800 | 0 | -5.95188 | -7.79385 | -4.10992 |
| | | | | | | | | | | | day 1 - day 5 | -14.8148 | 800 | 0 | -7.26948 | -9.50213 | -5.03682 |
| | | | | | | | | | | | day 2 - day 3 | -3.83333 | 800 | 0 | -1.88098 | -2.57307 | -1.18888 |
| | | | | | | | | | | | day 2 - day 4 | -6.33333 | 800 | 0 | -3.1077 | -4.12663 | -2.08877 |
| | | | | | | | | | | | day 2 - day 5 | -9.01852 | 800 | 0 | -4.42529 | -5.82001 | -3.03058 |
| | | | | | | | | | | | day 3 - day 4 | -2.5 | 800 | 0 | -1.22672 | -1.7682 | -0.68524 |
| | | | | | | | | | | | day 3 - day 5 | -5.18519 | 800 | 0 | -2.54432 | -3.40875 | -1.67988 |
| | | | | | | | | | | | day 4 - day 5 | -2.68519 | 800 | 0 | -1.31759 | -1.87824 | -0.75694 |
| | | | | | | | | | | Middle | day 1 - day 2 | -3.03333 | 800 | 0 | -1.48843 | -2.07371 | -0.90314 |
| | | | | | | | | | | | day 1 - day 3 | -5.03333 | 800 | 0 | -2.4698 | -3.30499 | -1.63462 |
| | | | | | | | | | | | day 1 - day 4 | -7.28333 | 800 | 0 | -3.57386 | -4.71722 | -2.43049 |
| | | | | | | | | | | | day 1 - day 5 | -8.73333 | 800 | 0 | -4.28536 | -5.63379 | -2.93693 |
| | | | | | | | | | | | day 2 - day 3 | -2 | 800 | 9.91E-07 | -0.98138 | -1.45901 | -0.50375 |
| | | | | | | | | | | | day 2 - day 4 | -4.25 | 800 | 0 | -2.08543 | -2.8186 | -1.35226 |
| | | | | | | | | | | | day 2 - day 5 | -5.7 | 800 | 0 | -2.79693 | -3.72158 | -1.87228 |
| | | | | | | | | | | | day 3 - day 4 | -2.25 | 800 | 1.51E-08 | -1.10405 | -1.60555 | -0.60255 |
| | | | | | | | | | | | day 3 - day 5 | -3.7 | 800 | 0 | -1.81555 | -2.47988 | -1.15123 |
| | | | | | | | | | | | day 4 - day 5 | -1.45 | 800 | 0.000998 | -0.7115 | -1.14321 | -0.27979 |
| | | | | | | | | | | Older | day 1 - day 2 | -2.14815 | 800 | 5.69E-07 | -1.05407 | -1.56124 | -0.54691 |
| | | | | | | | | | | | day 1 - day 3 | -3.48148 | 800 | 0 | -1.70833 | -2.35823 | -1.05843 |
| | | | | | | | | | | | day 1 - day 4 | -4.83333 | 800 | 0 | -2.37167 | -3.19001 | -1.55332 |
| | | | | | | | | | | | day 1 - day 5 | -6 | 800 | 0 | -2.94414 | -3.91767 | -1.9706 |
| | | | | | | | | | | | day 2 - day 3 | -1.33333 | 800 | 0.006351 | -0.65425 | -1.09562 | -0.21289 |
| | | | | | | | | | | | day 2 - day 4 | -2.68519 | 800 | 0 | -1.31759 | -1.87824 | -0.75694 |
| | | | | | | | | | | | day 2 - day 5 | -3.85185 | 800 | 0 | -1.89006 | -2.58442 | -1.19571 |
| | | | | | | | | | | | day 3 - day 4 | -1.35185 | 800 | 0.005387 | -0.66334 | -1.10594 | -0.22074 |
| | | | | | | | | | | | day 3 - day 5 | -2.51852 | 800 | 0 | -1.23581 | -1.77918 | -0.69245 |
| | | | | | | | | | | | day 4 - day 5 | -1.16667 | 800 | 0.0251 | -0.57247 | -1.00332 | -0.14162 |
| Accuracy | LMER | Per. Correct | AGE,DAY | (1 + 1\|ID) | AGE | 2 | 24.99996 | 6.075138 | 0.007075 | day 1 | Young - Middle | -0.06364 | 35.37045 | 0.27931 | -0.6517 | -1.52556 | 0.222155 |
| | | | | | DAY | 4 | 800 | 17.144 | 1.68E-13 | | Young - Older | 0.124837 | 35.37045 | 0.014414 | 1.278457 | 0.324453 | 2.232462 |
| | | | | | AGE:DAY | 8 | 800 | 4.508314 | 2.25E-05 | | Middle - Older | 0.188473 | 35.37045 | 0.000153 | 1.930162 | 0.901015 | 2.959308 |
| | | | | | | | | | | day 2 | Young - Middle | -0.03252 | 35.37045 | 0.709508 | -0.33301 | -1.19064 | 0.524622 |
| | | | | | | | | | | | Young - Older | 0.14517 | 35.37045 | 0.004066 | 1.486695 | 0.506044 | 2.467345 |
| | | | | | | | | | | | Middle - Older | 0.177687 | 35.37045 | 0.000332 | 1.819706 | 0.808736 | 2.830676 |
| | | | | | | | | | | day 3 | Young - Middle | -0.01486 | 35.37045 | 0.930278 | -0.15215 | -1.00519 | 0.700889 |
| | | | | | | | | | | | Young - Older | 0.061324 | 35.37045 | 0.322883 | 0.628022 | -0.26591 | 1.52195 |
| | | | | | | | | | | | Middle - Older | 0.076181 | 35.37045 | 0.165728 | 0.780174 | -0.10306 | 1.663407 |
| | | | | | | | | | | day 4 | Young - Middle | -0.02676 | 35.37045 | 0.791957 | -0.27405 | -1.12982 | 0.581711 |
| | | | | | | | | | | | Young - Older | 0.080799 | 35.37045 | 0.147543 | 0.827461 | -0.08089 | 1.735809 |
| | | | | | | | | | | | Middle - Older | 0.107559 | 35.37045 | 0.033275 | 1.101514 | 0.18815 | 2.014877 |
| | | | | | | | | | | day 5 | Young - Middle | -0.01066 | 35.37045 | 0.963459 | -0.10914 | -0.96159 | 0.743315 |
| | | | | | | | | | | | Young - Older | 0.071427 | 35.37045 | 0.219854 | 0.731492 | -0.16945 | 1.632436 |
| | | | | | | | | | | | Middle - Older | 0.082084 | 35.37045 | 0.126358 | 0.840628 | -0.04756 | 1.728812 |
| | | | | | | | | | | Young | day 1 - day 2 | -0.04374 | 800 | 0.13722 | -0.44794 | -0.86085 | -0.03503 |
| | | | | | | | | | | | day 1 - day 3 | -0.05876 | 800 | 0.015702 | -0.60173 | -1.03178 | -0.17168 |
| | | | | | | | | | | | day 1 - day 4 | -0.05397 | 800 | 0.033935 | -0.55275 | -0.97687 | -0.12862 |
| | | | | | | | | | | | day 1 - day 5 | -0.07742 | 800 | 0.000402 | -0.79284 | -1.24979 | -0.33589 |
| | | | | | | | | | | | day 2 - day 3 | -0.01502 | 800 | 0.930852 | -0.15379 | -0.54704 | 0.239457 |
| | | | | | | | | | | | day 2 - day 4 | -0.01023 | 800 | 0.982589 | -0.10481 | -0.49661 | 0.286998 |
| | | | | | | | | | | | day 2 - day 5 | -0.03368 | 800 | 0.378822 | -0.3449 | -0.74885 | 0.059053 |
| | | | | | | | | | | | day 3 - day 4 | 0.004783 | 800 | 0.999075 | 0.048985 | -0.34184 | 0.439809 |
| | | | | | | | | | | | day 3 - day 5 | -0.01866 | 800 | 0.858529 | -0.19111 | -0.58582 | 0.203605 |
| | | | | | | | | | | | day 4 - day 5 | -0.02344 | 800 | 0.723353 | -0.24009 | -0.63719 | 0.157008 |
| | | | | | | | | | | Middle | day 1 - day 2 | -0.01262 | 800 | 0.954707 | -0.12925 | -0.50177 | 0.243273 |
| | | | | | | | | | | | day 1 - day 3 | -0.00998 | 800 | 0.980728 | -0.10218 | -0.47395 | 0.269585 |
| | | | | | | | | | | | day 1 - day 4 | -0.0171 | 800 | 0.873316 | -0.1751 | -0.54929 | 0.199097 |
| | | | | | | | | | | | day 1 - day 5 | -0.02444 | 800 | 0.646622 | -0.25027 | -0.62827 | 0.127727 |
| | | | | | | | | | | | day 2 - day 3 | 0.002643 | 800 | 0.999891 | 0.027066 | -0.34353 | 0.397661 |
| | | | | | | | | | | | day 2 - day 4 | -0.00448 | 800 | 0.999123 | -0.04585 | -0.41661 | 0.324912 |
| | | | | | | | | | | | day 2 - day 5 | -0.01182 | 800 | 0.964192 | -0.12102 | -0.4933 | 0.251247 |
| | | | | | | | | | | | day 3 - day 4 | -0.00712 | 800 | 0.994636 | -0.07291 | -0.44406 | 0.298234 |
| | | | | | | | | | | | day 3 - day 5 | -0.01446 | 800 | 0.927235 | -0.14809 | -0.52124 | 0.225056 |
| | | | | | | | | | | | day 4 - day 5 | -0.00734 | 800 | 0.993967 | -0.07518 | -0.44637 | 0.296012 |
| | | | | | | | | | | Older | day 1 - day 2 | -0.02341 | 800 | 0.724568 | -0.2397 | -0.63678 | 0.157377 |
| | | | | | | | | | | | day 1 - day 3 | -0.12227 | 800 | 0 | -1.25217 | -1.79337 | -0.71097 |
| | | | | | | | | | | | day 1 - day 4 | -0.09801 | 800 | 2.31E-06 | -1.00374 | -1.49642 | -0.51107 |
| | | | | | | | | | | | day 1 - day 5 | -0.13083 | 800 | 0 | -1.33981 | -1.89948 | -0.78013 |
| | | | | | | | | | | | day 2 - day 3 | -0.09886 | 800 | 1.82E-06 | -1.01247 | -1.50673 | -0.5182 |
| | | | | | | | | | | | day 2 - day 4 | -0.07461 | 800 | 0.000745 | -0.76404 | -1.21658 | -0.3115 |

| | | | | | | | | | | | | | | | | | |
|---|---|---|---|---|---|---|---|---|---|---|---|---|---|---|---|---|---|
| | | | | | | | | | | | day 2 - day 5 | -0.10742 | 800 | 1.46E-07 | -1.1001 | -1.61086 | -0.58934 |
| | | | | | | | | | | | day 3 - day 4 | 0.024258 | 800 | 0.696966 | 0.248425 | -0.14913 | 0.645984 |
| | | | | | | | | | | | day 3 - day 5 | -0.00856 | 800 | 0.991132 | -0.08764 | -0.47906 | 0.30379 |
| | | | | | | | | | | | day 4 - day 5 | -0.03282 | 800 | 0.406178 | -0.33606 | -0.73935 | 0.067223 |
| | | | | | AGE | 2 | 25 | 12.07554 | 0.000214 | | Young - Middle | 7.681741 | 31.64578 | 0.015768 | 1.911783 | 0.473198 | 3.350368 |
| | | | | | DAY | 2 | 218 | 10.40153 | 4.85E-05 | day 5 | Young - Older | 11.81251 | 31.64578 | 0.000305 | 2.939823 | 1.323855 | 4.555791 |
| | | | | | AGE:DAY | 4 | 218 | 1.377101 | 0.24284 | | Middle - Older | 4.130771 | 31.64578 | 0.265335 | 1.02804 | -0.32658 | 2.382655 |
| | | | | | | | | | | | Young - Middle | 8.009999 | 31.64578 | 0.011538 | 1.993478 | 0.544917 | 3.442039 |
| | | | | | | | | | | day 10 | Young - Older | 12.22752 | 31.64578 | 0.000196 | 3.043107 | 1.409969 | 4.676245 |
| | | | | | | | | | | | Middle - Older | 4.217519 | 31.64578 | 0.25147 | 1.049629 | -0.30648 | 2.405739 |
| | | | | | | | | | | | Young - Middle | 10.30596 | 31.64578 | 0.001119 | 2.564881 | 1.037311 | 4.092451 |
| FU Scores | LMER | Score | AGE,DAY | (1 + 1\|ID) | | | | | | day 60 | Young - Older | 11.73988 | 31.64578 | 0.00033 | 2.921747 | 1.308741 | 4.534753 |
| | | | | | | | | | | | Middle - Older | 1.433923 | 31.64578 | 0.84652 | 0.356866 | -0.96641 | 1.680145 |
| | | | | | | | | | | | day 5 - day 10 | -0.60415 | 218 | 0.845334 | -0.15036 | -0.70682 | 0.406107 |
| | | | | | | | | | | Young | day 5 - day 60 | 1.403806 | 218 | 0.405902 | 0.34937 | -0.2151 | 0.913843 |
| | | | | | | | | | | | day 10 - day 60 | 2.007953 | 218 | 0.160307 | 0.499727 | -0.07486 | 1.074318 |
| | | | | | | | | | | | day 5 - day 10 | -0.27589 | 218 | 0.961775 | -0.06866 | -0.59523 | 0.457908 |
| | | | | | | | | | | Middle | day 5 - day 60 | 4.028022 | 218 | 0.000402 | 1.002468 | 0.396191 | 1.608746 |
| | | | | | | | | | | | day 10 - day 60 | 4.303911 | 218 | 0.000142 | 1.07113 | 0.454343 | 1.687917 |
| | | | | | | | | | | | day 5 - day 10 | -0.18914 | 218 | 0.983646 | -0.04707 | -0.60188 | 0.507735 |
| | | | | | | | | | | Older | day 5 - day 60 | 1.331174 | 218 | 0.444252 | 0.331294 | -0.23219 | 0.894782 |
| | | | | | | | | | | | day 10 - day 60 | 1.520315 | 218 | 0.347788 | 0.378366 | -0.18779 | 0.944524 |
| | | | | | AGE | 2 | 25 | 8.715846 | 0.001343 | | | | | | | | |
| Rand. Blocks | LMER | Score | AGE,DAY | (1 + 1\|ID) | DAY | 5 | 125 | 2.02342 | 0.079779 | | | | | | | | |
| | | | | | AGE:DAY | 10 | 125 | 0.783287 | 0.644731 | | | | | | | | |
| SICIrest at Baseline | LM | SICI ratio (SICI/Test) | AGE | NA | AGE | 2 | | 0.097957 | 0.907064 | | | | | | | | |
| | | | | | AGE | 2 | | 0.268557 | 0.765672 | | | | | | | | |
| SICIrest change Tr. D1 | LM | ΔSICI ratio | AGE,TimePoint | NA | TimePoint | 1 | | 0.02577 | 0.873166 | | | | | | | | |
| | | | | | AGE:TimePoint | 2 | | 0.111119 | 0.895071 | | | | | | | | |
| | | | | | AGE | 2 | | 0.034383 | 0.966226 | | | | | | | | |
| SICIrest change Tr. Week | LM | ΔSICI ratio | AGE,TimePoint | NA | TimePoint | 1 | | 0.803903 | 0.374499 | | | | | | | | |
| | | | | | AGE:TimePoint | 2 | | 0.244467 | 0.784112 | | | | | | | | |

**Table 3. Statistical tests run on data from the second experiment, comparing verum and placebo groups for each age group.**

| | | | | | | ANOVA | | | | | PostHoc tests | | | | | | | |
|---|---|---|---|---|---|---|---|---|---|---|---|---|---|---|---|---|---|---|
| Aspect | Age Group | Model | Dependent | Independent | Random | ANOVA param. | DF num. | DF den. | F | p | Level | Contrast | Estimate | DF | p | d | CI | |
| Baseline | Young | LM | Score | STIM | NA | STIM | 1 | | 0.52376 | 0.479083 | | | | | | | | |
| | Middle | | | | | | 1 | | 1.849072 | 0.191654 | | | | | | | | |
| | Older | | | | | | 1 | | 0.015125 | 0.903289 | | | | | | | | |
| Block 1 Accuracy | Young | LM | Per. Correct | STIM | NA | STIM | 1 | | 1.229327 | 0.282978 | | | | | | | | |
| | Middle | | | | | | 1 | | 2.850293 | 0.109615 | | | | | | | | |
| | Older | | | | | | 1 | | 0.038756 | 0.845826 | | | | | | | | |
| Training | Young | LMER | Score | STIM,DAY | (1 + 1\|ID) | STIM | 1 | 17 | 0.090493 | 0.767199 | day 1 | Verum vs. Placebo | 0.582313 | 20.04746 | 0.84647 | 0.116902 | -1.12665 | 1.360452 |
| | | | | | | DAY | 4 | 543 | 151.4861 | 6.11E-87 | day 2 | | 1.5869 | 20.04746 | 0.598851 | 0.318578 | -0.92831 | 1.565463 |
| | | | | | | STIM:DAY | 4 | 543 | 5.150391 | 0.000444 | day 3 | | 2.201949 | 20.04746 | 0.466866 | 0.442052 | -0.80839 | 1.692493 |
| | | | | | | | | | | | day 4 | | 2.597604 | 20.04746 | 0.391947 | 0.521482 | -0.73185 | 1.774814 |
| | | | | | | | | | | | day 5 | | -2.68403 | 20.04746 | 0.376683 | -0.53883 | -1.79286 | 0.715193 |
| | Middle | LMER | Score | STIM,DAY | (1 + 1\|ID) | STIM | 1 | 17 | 0.027385 | 0.870515 | day 1 | Verum vs. Placebo | -1.45057 | 20.31419 | 0.483438 | -0.41001 | -1.61351 | 0.793501 |
| | | | | | | DAY | 4 | 543 | 140.4404 | 2.48E-82 | day 2 | | -0.36435 | 20.31419 | 0.859477 | -0.10298 | -1.3003 | 1.094332 |
| | | | | | | STIM:DAY | 4 | 543 | 3.380069 | 0.009564 | day 3 | | 1.051544 | 20.31419 | 0.610406 | 0.297221 | -0.90315 | 1.497596 |
| | | | | | | | | | | | day 4 | | 0.787435 | 20.31419 | 0.702407 | 0.22257 | -0.97628 | 1.421418 |
| | | | | | | | | | | | day 5 | | 1.583947 | 20.31419 | 0.444677 | 0.447705 | -0.75707 | 1.652481 |
| | Older | LMER | Score | STIM,DAY | (1 + 1\|ID) | STIM | 1 | 20.99994 | 4.997128 | 0.03638 | day 1 | Verum vs. Placebo | 2.771352 | 24.74045 | 0.105118 | 0.934891 | -0.24504 | 2.114822 |
| | | | | | | DAY | 4 | 659 | 95.52916 | 4.39E-64 | day 2 | | 4.138842 | 24.74045 | 0.018897 | 1.396202 | 0.174931 | 2.617473 |
| | | | | | | STIM:DAY | 4 | 659 | 2.719338 | 0.028847 | day 3 | | 2.583184 | 24.74045 | 0.129596 | 0.871414 | -0.30402 | 2.046852 |
| | | | | | | | | | | | day 4 | | 3.622439 | 24.74045 | 0.03747 | 1.221998 | 0.018087 | 2.425908 |
| | | | | | | | | | | | day 5 | | 4.558031 | 24.74045 | 0.010556 | 1.537611 | 0.300761 | 2.774462 |
| Online Learning D1 | Young | LM | ΔScore | STIM | NA | STIM | 1 | | 0.005869 | 0.93983 | | | | | | | | |
| | Middle | LM | ΔScore | STIM | NA | STIM | 1 | | 0.091144 | 0.766389 | | | | | | | | |
| | Older | LM | ΔScore | STIM | NA | STIM | 1 | | 9.192547 | 0.006341 | day 1 | Verum vs. Placebo | 3.619165 | 21 | 0.006341 | 1.295378 | 0.322132 | 2.268624 |
| Online learning D2-D5 | Young | LMER | ΔScore | STIM,DAY | (1 + 1\|ID) | STIM | 1 | 17 | 0.170196 | 0.685094 | | | | | | | | |
| | | | | | | DAY | 3 | 51 | 1.493668 | 0.227352 | | | | | | | | |
| | | | | | | STIM:DAY | 3 | 51 | 0.228043 | 0.876449 | | | | | | | | |
| | Middle | LM | ΔScore | STIM,DAY | NA | STIM | 1 | | 0.130912 | 0.71861 | | | | | | | | |
| | | | | | | DAY | 3 | | 3.70962 | 0.015589 | | | | | | | | |
| | | | | | | STIM:DAY | 3 | | 0.282582 | 0.837797 | | | | | | | | |
| | Older | LMER | ΔScore | STIM,DAY | (1 + 1\|ID) | STIM | 1 | 21 | 0.007162 | 0.933356 | | | | | | | | |
| | | | | | | DAY | 3 | 63 | 1.337383 | 0.270225 | | | | | | | | |
| | | | | | | STIM:DAY | 3 | 63 | 0.504896 | 0.6803 | | | | | | | | |
| Online slope D1 | Young | LM | Slope | STIM | NA | STIM | 1 | | 0.007103 | 0.933821 | | | | | | | | |
| | Middle | LM | Slope | STIM | NA | STIM | 1 | | 0.226887 | 0.639907 | | | | | | | | |
| | Older | LM | Slope | STIM | NA | STIM | 1 | | 7.228349 | 0.013754 | day 1 | Verum vs. Placebo | 3.499548 | 21 | 0.013754 | 1.148678 | 0.192907 | 2.104449 |
| Online slope D2-D5 | Young | LMER | Slope | STIM,DAY | (1 + 1\|ID) | STIM | 1 | 17 | 0.001108 | 0.973835 | | | | | | | | |
| | | | | | | DAY | 3 | 51 | 2.221858 | 0.096831 | | | | | | | | |
| | | | | | | STIM:DAY | 3 | 51 | 0.61681 | 0.607289 | | | | | | | | |
| | Middle | LM | Slope | STIM,DAY | NA | STIM | 1 | | 1.038524 | 0.311777 | | | | | | | | |
| | | | | | | DAY | 3 | | 7.939899 | 0.000129 | | | | | | | | |
| | | | | | | STIM:DAY | 3 | | 1.413525 | 0.246378 | | | | | | | | |
| | Older | LMER | Slope | STIM,DAY | (1 + 1\|ID) | STIM | 1 | 21.00008 | 0.229233 | 0.637041 | | | | | | | | |
| | | | Slope | | | DAY | 3 | 62.99986 | 1.757451 | 0.16441 | | | | | | | | |
| | | | Slope | | | STIM:DAY | 3 | 62.99986 | 0.49405 | 0.687702 | | | | | | | | |
| Offline Learning | Young | LM | ΔScore | STIM,NIGHT | NA | STIM | 1 | | 0.0229 | 0.880166 | | | | | | | | |
| | | | | | | NIGHT | 3 | | 0.147828 | 0.930739 | | | | | | | | |
| | | | | | | STIM:NIGHT | 3 | | 0.447888 | 0.719593 | | | | | | | | |
| | Middle | LM | ΔScore | STIM,NIGHT | NA | STIM | 1 | | 0.210953 | 0.647487 | | | | | | | | |
| | | | | | | NIGHT | 3 | | 1.281362 | 0.2878 | | | | | | | | |
| | | | | | | STIM:NIGHT | 3 | | 0.125095 | 0.944954 | | | | | | | | |
| | Older | LM | ΔScore | STIM,NIGHT | NA | STIM | 1 | | 0.040191 | 0.841592 | | | | | | | | |
| | | | | | | NIGHT | 3 | | 1.314234 | 0.275203 | | | | | | | | |
| | | | | | | STIM:NIGHT | 3 | | 1.630893 | 0.188352 | | | | | | | | |
| Speed | Young | LMER | Seq. Number | STIM,DAY | (1 + 1\|ID) | STIM | 1 | 17 | 2.078396 | 0.167566 | day 1 | Verum vs. Placebo | 1.416667 | 18.36444 | 0.59824 | 0.462183 | -1.35172 | 2.276081 |
| | | | | | | DAY | 4 | 543 | 511.9617 | 1.21E-182 | day 2 | | 3.103704 | 18.36444 | 0.255089 | 1.012572 | -0.82261 | 2.847754 |
| | | | | | | STIM:DAY | 4 | 543 | 6.612859 | 3.35E-05 | day 3 | | 4.52037 | 18.36444 | 0.103918 | 1.474755 | -0.39015 | 3.339664 |
| | | | | | | | | | | | day 4 | | 4.787037 | 18.36444 | 0.08638 | 1.561754 | -0.30992 | 3.433428 |
| | | | | | | | | | | | day 5 | | 4.851852 | 18.36444 | 0.082529 | 1.5829 | -0.29047 | 3.456273 |
| | Middle | LMER | Seq. Number | STIM,DAY | (1 + 1\|ID) | STIM | 1 | 16.99999 | 2.2041 | 0.15595 | day 1 | Verum vs. Placebo | 0.948148 | 19.20977 | 0.547837 | 0.421402 | -1.02501 | 1.867819 |
| | | | | | | DAY | 4 | 543 | 330.4279 | 6.51E-144 | day 2 | | 1.896296 | 19.20977 | 0.235893 | 0.842805 | -0.62102 | 2.306629 |
| | | | | | | STIM:DAY | 4 | 543 | 3.795345 | 0.0047 | day 3 | | 2.859259 | 19.20977 | 0.080514 | 1.270791 | -0.22213 | 2.763708 |
| | | | | | | | | | | | day 4 | | 2.553704 | 19.20977 | 0.115649 | 1.134988 | -0.34749 | 2.617463 |
| | | | | | | | | | | | day 5 | | 2.9 | 19.20977 | 0.076617 | 1.288899 | -0.20549 | 2.783291 |
| | Older | LMER | Seq. Number | STIM,DAY | (1 + 1\|ID) | STIM | 1 | 21 | 0.496911 | 0.488606 | | | | | | | | |
| | | | | | | DAY | 4 | 659 | 264.3163 | 2.17E-135 | | | | | | | | |
| | | | | | | STIM:DAY | 4 | 659 | 0.703884 | 0.589453 | | | | | | | | |
| Accuracy | Young | LMER | Per. Correct | STIM,DAY | (1 + 1\|ID) | STIM | 1 | 17 | 0.155273 | 0.698446 | day 1 | Verum vs. Placebo | 0.00426 | 22.85555 | 0.89813 | 0.058486 | -0.8766 | 0.993576 |
| | | | | | | DAY | 4 | 543 | 6.412699 | 4.78E-05 | day 2 | | -0.00041 | 22.85555 | 0.990247 | -0.00558 | -0.9405 | 0.929338 |
| | | | | | | STIM:DAY | 4 | 543 | 5.873906 | 0.000124 | day 3 | | 0.001119 | 22.85555 | 0.97316 | 0.015368 | -0.91956 | 0.950299 |
| | | | | | | | | | | | day 4 | | 0.005933 | 22.85555 | 0.858491 | 0.081466 | -0.85378 | 1.016716 |
| | | | | | | | | | | | day 5 | | -0.0711 | 22.85555 | 0.041439 | -0.97621 | -1.95743 | 0.00501 |
| | | | | | | | | | | | Verum | day 1 - day 2 | -0.03907 | 543 | 0.028267 | -0.53649 | -0.94824 | -0.12474 |
| | | | | | | | | | | | | day 2 - day 3 | -0.01654 | 543 | 0.725431 | -0.22714 | -0.61126 | 0.156976 |
| | | | | | | | | | | | | day 3 - day 4 | -3.08E-05 | 543 | 1 | -0.00042 | -0.37824 | 0.377393 |
| | | | | | | | | | | | | day 4 - day 5 | 0.053588 | 543 | 0.000608 | 0.73578 | 0.296299 | 1.175261 |
| | | | | | | | | | | | Placebo | day 1 - day 2 | -0.04374 | 543 | 0.016238 | -0.60056 | -1.03895 | -0.16217 |
| | | | | | | | | | | | | day 2 - day 3 | -0.01502 | 543 | 0.821188 | -0.20619 | -0.60938 | 0.196999 |
| | | | | | | | | | | | | day 3 - day 4 | 0.004783 | 543 | 0.99708 | 0.065675 | -0.33308 | 0.464432 |
| | | | | | | | | | | | | day 4 - day 5 | -0.02344 | 543 | 0.451833 | -0.3219 | -0.73208 | 0.088289 |
| | Middle | LMER | Per. Correct | STIM,DAY | (1 + 1\|ID) | STIM | 1 | 17 | 4.111138 | 0.058575 | day 1 | Verum vs. Placebo | -0.14785 | 21.71704 | 0.002655 | -1.67475 | -2.82044 | -0.52907 |
| | | | | | | DAY | 4 | 543 | 13.05669 | 3.69E-10 | day 2 | | -0.10713 | 21.71704 | 0.022432 | -1.21357 | -2.30346 | -0.12368 |
| | | | | | | STIM:DAY | 4 | 543 | 7.065989 | 1.50E-05 | day 3 | | -0.06873 | 21.71704 | 0.12927 | -0.77855 | -1.83055 | 0.273456 |
| | | | | | | | | | | | day 4 | | -0.04822 | 21.71704 | 0.280733 | -0.54616 | -1.58437 | 0.492054 |
| | | | | | | | | | | | day 5 | | -0.04364 | 21.71704 | 0.327711 | -0.49435 | -1.53013 | 0.541425 |
| | Older | LMER | Per. Correct | STIM,DAY | (1 + 1\|ID) | STIM | 1 | 21 | 9.092071 | 0.006587 | day 1 | Verum vs. Placebo | 0.113522 | 35.17942 | 0.00743 | 0.927315 | 0.20904 | 1.64559 |
| | | | | | | DAY | 4 | 659 | 3.554149 | 0.007023 | day 2 | | 0.171254 | 35.17942 | 0.000134 | 1.398903 | 0.615213 | 2.182592 |
| | | | | | | STIM:DAY | 4 | 659 | 4.054401 | 0.002959 | day 3 | | 0.0529 | 35.17942 | 0.194073 | 0.432117 | -0.24288 | 1.107118 |
| | | | | | | | | | | | day 4 | | 0.100713 | 35.17942 | 0.016405 | 0.822686 | 0.115919 | 1.529453 |
| | | | | | | | | | | | day 5 | | 0.09076 | 35.17942 | 0.029345 | 0.741386 | 0.042722 | 1.44005 |
| FU Scores | Young | LMER | Score | STIM,DAY | (1 + 1\|ID) | STIM | 1 | 16.99982 | 0.293404 | 0.595076 | day | day 5 - day 10 | -0.94675 | 148 | 0.558089 | -0.19353 | -0.58954 | 0.202492 |
| | | | | | | DAY | 2 | 148.0001 | 3.47365 | 0.033554 | | day 5 - day 60 | 1.454159 | 148 | 0.255451 | 0.297245 | -0.10481 | 0.699304 |
| | | | | | | STIM:DAY | 2 | 148.0001 | 0.10859 | 0.897169 | | day 10 - day 60 | 2.400907 | 148 | 0.02638 | 0.490771 | 0.071136 | 0.910406 |
| | Middle | LMER | Score | STIM,DAY | (1 + 1\|ID) | STIM | 1 | 17 | 0.706449 | 0.412291 | day | day 5 - day 10 | -1.54311 | 148 | 0.088768 | -0.39813 | -0.80832 | 0.012069 |
| | | | | | | DAY | 2 | 148 | 14.3154 | 2.07E-06 | | day 5 - day 60 | 2.321085 | 148 | 0.004877 | 0.598845 | 0.166169 | 1.03152 |
| | | | | | | STIM:DAY | 2 | 148 | 2.971993 | 0.054266 | | day 10 - day 60 | 3.864197 | 148 | 1.15E-06 | 0.996971 | 0.499612 | 1.49433 |
| | Older | LMER | Score | STIM,DAY | (1 + 1\|ID) | STIM | 1 | 20.99996 | 5.698807 | 0.026458 | stim | Verum vs. Placebo | 4.916899 | 21 | 0.026458 | 1.490656 | 0.113987 | 2.867324 |
| | | | | | | DAY | 2 | 180 | 5.581149 | 0.00445 | day | day 5 - day 10 | -0.43619 | 182 | 0.70441 | -0.13595 | -0.48853 | 0.21663 |
| | | | | | | STIM:DAY | 2 | 180 | 0.122867 | 0.884456 | | day 5 - day 60 | 1.409152 | 182 | 0.028644 | 0.439199 | 0.064565 | 0.813833 |
| | | | | | | | | | | | | day 10 - day 60 | 1.845337 | 182 | 0.002558 | 0.575147 | 0.183955 | 0.96634 |
| SICI at Baseline | Young | LM | SICI ratio (SICI/Test) | STIM | NA | STIM | 1 | | 1.209097 | 0.287781 | | | | | | | | |
| | Middle | LM | SICI ratio (SICI/Test) | STIM | NA | STIM | 1 | | 1.625375 | 0.221739 | | | | | | | | |
| | Older | LM | SICI ratio (SICI/Test) | STIM | NA | STIM | 1 | | 0.076384 | 0.78603 | | | | | | | | |
| SICIrest change Tr. D1 | Young | LM | ΔSICI ratio | STIM,TimePoint | NA | STIM | 1 | | 2.838527 | 0.101759 | | | | | | | | |
| | | | | | | TimePoint | 1 | | 0.251919 | 0.619163 | | | | | | | | |
| | | | | | | STIM:TimePoint | 1 | | 0.016824 | 0.897611 | | | | | | | | |
| | Middle | LM | ΔSICI ratio | STIM,TimePoint | NA | STIM | 1 | | 2.457586 | 0.127807 | | | | | | | | |
| | | | | | | TimePoint | 1 | | 0.039016 | 0.844795 | | | | | | | | |
| | | | | | | STIM:TimePoint | 1 | | 0.007487 | 0.931641 | | | | | | | | |
| | Older | LM | ΔSICI ratio | STIM,TimePoint | NA | STIM | 1 | | 0.016204 | 0.899556 | | | | | | | | |
| | | | | | | TimePoint | 1 | | 0.002271 | 0.962305 | | | | | | | | |
| | | | | | | STIM:TimePoint | 1 | | 0.2572 | 0.615758 | | | | | | | | |
| SICIrest change Tr. Week | Young | LM | ΔSICI ratio | STIM,TimePoint | NA | STIM | 1 | | 0.979849 | 0.329662 | | | | | | | | |
| | | | | | | TimePoint | 1 | | 0.14306 | 0.707755 | | | | | | | | |
| | | | | | | STIM:TimePoint | 1 | | 0.111089 | 0.741082 | | | | | | | | |
| | Middle | LM | ΔSICI ratio | STIM,TimePoint | NA | STIM | 1 | | 0.895513 | 0.351306 | | | | | | | | |
| | | | | | | TimePoint | 1 | | 0.080159 | 0.778965 | | | | | | | | |
| | | | | | | STIM:TimePoint | 1 | | 0.692909 | 0.411544 | | | | | | | | |
| | Older | LM | ΔSICI ratio | STIM,TimePoint | NA | STIM | 1 | | 0.027865 | 0.868548 | | | | | | | | |
| | | | | | | TimePoint | 1 | | 0.45041 | 0.507275 | | | | | | | | |
| | | | | | | STIM:TimePoint | 1 | | 0.018168 | 0.89368 | | | | | | | | |

**Table 4. Statistical tests run on data from the second experiment, comparing young-like and old-like older adults in the verum group. Labels are either "Young-like" or "Old-Like"**

| | | | | | ANOVA | | | | | PostHoc tests | | | | | | | |
|---|---|---|---|---|---|---|---|---|---|---|---|---|---|---|---|---|---|
| **Aspect** | **Model** | **Dependent** | **Independent** | **Random** | **ANOVA param.** | **DF num.** | **DF den.** | **F** | **p** | **Level** | **Contrast** | **Estimate** | **DF** | **p** | **d** | **CI** | |
| Baseline Speed | LM | Seq. Number | LABEL | NA | LABEL | 1 | | 6.0615 | 0.029928 | BL | YoungLike - OldLike | 4.208333 | 12 | 0.029928 | 1.329638 | 0.077816 | 2.581461 |
| Block 1 Speed | LM | Seq. Number | LABEL | NA | LABEL | 1 | | 6.817824 | 0.022756 | B1 | YoungLike - OldLike | 4.25 | 12 | 0.022756 | 1.410153 | 0.149271 | 2.671036 |
| Block 1 Accuracy | LM | Per. Correct | LABEL | NA | LABEL | 1 | | 0.097421 | 0.760304 | | | | | | | | |
| Training | LMER | Score | LABEL,DAY | (1 + 1\|ID) | LABEL | 1 | 12 | 23.68295 | 0.000387 | day 1 | YoungLike - OldLike | 3.896406 | 16.15486 | 0.020478 | 1.361515 | 0.16111 | 2.56192 |
| | | | | | DAY | 4 | 398 | 84.58827 | 6.20E-52 | day 2 | | 6.578695 | 16.15486 | 0.000499 | 2.298783 | 0.966261 | 3.631304 |
| | | | | | LABEL:DAY | 4 | 398 | 9.701391 | 1.70E-07 | day 3 | | 7.004135 | 16.15486 | 0.000278 | 2.447443 | 1.089343 | 3.805544 |
| | | | | | | | | | | day 4 | | 7.411037 | 16.15486 | 0.00016 | 2.589626 | 1.206045 | 3.973207 |
| | | | | | | | | | | day 5 | | 9.362692 | 16.15486 | 1.28E-05 | 3.27159 | 1.753634 | 4.789546 |
| Online slope D1 | LM | Slope | LABEL,BLOCK | NA | LABEL | 1 | | 35.0624 | 7.62E-08 | | | | | | | | |
| | | | | | BLOCK | 1 | | 27.45138 | 1.28E-06 | | | | | | | | |
| | | | | | LABEL:BLOCK | 1 | | 2.214154 | 0.140681 | | | | | | | | |
| Online slope D2-D5 | LM | Slope | LABEL,BLOCK | NA | LABEL | 1 | | 289.9642 | 3.54E-47 | | | | | | | | |
| | | | | | BLOCK | 1 | | 13.87193 | 0.00023 | | | | | | | | |
| | | | | | LABEL:BLOCK | 1 | | 0.083031 | 0.773411 | | | | | | | | |
| Speed | LMER | Seq. Number | LABEL,DAY | (1 + 1\|ID) | DAY | 4 | 398 | 224.2692 | 1.49E-100 | day 1 | YoungLike - OldLike | 2.777778 | 13.2841 | 0.073675 | 1.719401 | -0.26609 | 3.704889 |
| | | | | | LABEL | 1 | 11.99999 | 16.75407 | 0.00149 | day 2 | | 5.590278 | 13.2841 | 0.001731 | 3.460294 | 1.257319 | 5.663269 |
| | | | | | DAY:LABEL | 4 | 398 | 26.0044 | 3.62E-19 | day 3 | | 6.180556 | 13.2841 | 0.000792 | 3.825667 | 1.562478 | 6.088856 |
| | | | | | | | | | | day 4 | | 6.270833 | 13.2841 | 0.000704 | 3.881547 | 1.608767 | 6.154328 |
| | | | | | | | | | | day 5 | | 7.722222 | 13.2841 | 0.000113 | 4.779934 | 2.340342 | 7.219527 |
| Speed slope D1 | LM | Seq. Number | LABEL,BLOCK | NA | LABEL | 1 | | 44.78938 | 2.75E-09 | Slope D1 | YoungLike - OldLike | 0.547619 | 80 | 0.005555 | 0.290895 | 0.070579 | 0.511211 |
| | | | | | BLOCK | 1 | | 27.93586 | 1.06E-06 | | | | | | | | |
| | | | | | LABEL:BLOCK | 1 | | 8.123515 | 0.005555 | | | | | | | | |
| Speed slope D2-D5 | LM | Seq. Number | LABEL,BLOCK | NA | LABEL | 1 | | 305.058 | 6.53E-49 | | | | | | | | |
| | | | | | BLOCK | 1 | | 13.34315 | 0.000301 | | | | | | | | |
| | | | | | LABEL:BLOCK | 1 | | 1.296887 | 0.255604 | | | | | | | | |
| Accuracy | LMER | Per. Correct | LABEL,Day | (1 + 1\|ID) | LABEL | 1 | 12 | 0.007673 | 0.931642 | Day | day 1 - day 2 | -0.08039 | 398 | 6.10E-05 | -0.71397 | -1.10482 | -0.32313 |
| | | | | | DAY | 4 | 398 | 11.02619 | 1.72E-08 | | day 1 - day 3 | -0.05978 | 398 | 0.0065 | -0.53093 | -0.89339 | -0.16847 |
| | | | | | LABEL:DAY | 4 | 398 | 0.305363 | 0.874357 | | day 1 - day 4 | -0.08503 | 398 | 1.80E-05 | -0.75521 | -1.15326 | -0.35715 |
| | | | | | | | | | | | day 1 - day 5 | -0.10883 | 398 | 1.41E-08 | -0.96656 | -1.40542 | -0.5277 |
| | | | | | | | | | | | day 2 - day 3 | 0.020609 | 398 | 0.766217 | 0.183043 | -0.14569 | 0.511781 |
| | | | | | | | | | | | day 2 - day 4 | -0.00464 | 398 | 0.998922 | -0.04124 | -0.3654 | 0.28293 |
| | | | | | | | | | | | day 2 - day 5 | -0.02844 | 398 | 0.485326 | -0.25259 | -0.58562 | 0.080446 |
| | | | | | | | | | | | day 3 - day 4 | -0.02525 | 398 | 0.603055 | -0.22428 | -0.55541 | 0.106847 |
| | | | | | | | | | | | day 3 - day 5 | -0.04905 | 398 | 0.043143 | -0.43563 | -0.78596 | -0.0853 |
| | | | | | | | | | | | day 4 - day 5 | -0.0238 | 398 | 0.656414 | -0.21135 | -0.54168 | 0.118976 |
